# Furan fatty acid supplementation protects against muscle atrophy during cancer cachexia

**DOI:** 10.64898/2026.09.08.750107

**Authors:** Anaïs Deglos, Jennifer Falconi, Charles Géminard, Christelle Bertrand-Gaday, Béatrice Bonafos, Vanessa Bauer, Lisa Heron-Milhavet, Erwann Durand, Pierre Delobel, Céline Jahannault-Talignani, Cécile Déjou, Anna Carval, Jatuporn Chaiyut, Laurent Vaysse, Siriluck Liengprayoon, Laurence Pessemesse, Charles Coudray, Walter Vetter, Christine Feillet-Coudray, Alexandre Djiane, François Casas

**Affiliations:** DMEM, Univ Montpellier, INRAE, Montpellier, France; IRCM, Univ Montpellier, INSERM, Montpellier, France; Institute of Food Chemistry (170b), University of Hohenheim, Stuttgart, Germany; Qualisud, Univ Montpellier, CIRAD, Montpellier, France; Qualisud, Univ Montpellier, Avignon Université, CIRAD, Institut Agro, IRD, Université de La Réunion, Montpellier, France; Kasetsart agricultural and agro-industrial product improvement institute, Kasetsart University, Bangkok, Thaïland; CIRAD, UPR BioWooEB, Montpellier, France; BioWooEB, Univ Montpellier, CIRAD, Montpellier, France

**Keywords:** Furan fatty acid, skeletal muscle, cancer cachexia, C26 mouse

## Abstract

**Background:** Cachexia is a multifactorial syndrome frequently observed in cancer patients, characterized by progressive weight loss, muscle atrophy, and systemic inflammation. We recently demonstrated that supplementation with FuFA-F2, a naturally occurring lipid found in various foods, increases muscle mass in different metabolic contexts. Here, we investigated whether FuFA-F2 supplementation could prevent tumor-induced muscle wasting and preserve skeletal muscle integrity during cancer cachexia.

**Methods:** *In vitro*, C2C12 myotubes were exposed to TNFα and IFNγ to mimic cachectic conditions, and the effects of FuFA-F2 on myotube morphology were assessed. *In vivo*, cancer cachexia was induced by subcutaneous injection of C26 adenocarcinoma cells into male CD2F1 mice. Three groups were compared: non-grafted control mice, untreated C26 tumor-bearing mice, and C26 tumor-bearing mice orally supplemented with FuFA-F2 (13 mg/kg/day) for 14 days.

**Results:** In C2C12 myotubes, TNFα and IFNγ exposure reduced myotube area by 17% (p < 0.05), whereas FuFA-F2 treatment prevented this atrophy and restored myotube area to control levels (p < 0.05). *In vivo*, FuFA-F2 supplementation prevented muscle wasting in C26 tumor-bearing mice without affecting tumor growth or the loss of white adipose tissue. After 14 days, hindlimb muscle weight was reduced by 22% in C26 mice compared with controls (0.706 vs. 0.903 g, p < 0.05), whereas muscle weight in FuFA-F2-treated mice (0.835 g) was not significantly different from controls. Consistently, spontaneous wheel activity was markedly reduced in C26 mice during the final four days (-79%; 3.4 vs. 16.5 km, p < 0.05), whereas FuFA-F2- treated mice maintained activity levels closer to those of controls (11.2 km). RNA-seq analysis revealed extensive transcriptional reprogramming of skeletal muscle in response to C26 tumor growth, with 5,465 differentially expressed genes (DEGs; 32% of detected genes) between Control and C26 mice. Notably, FuFA- F2 substantially attenuated this response, with only 366 DEGs (2%) between Control and C26 + FuFA-F2 mice, and principal component analysis (PCA) showed a transcriptomic profile closer to controls. Tumor-induced alterations involved pathways related to proteostasis, inflammation, and tissue remodeling, which were largely prevented or attenuated by FuFA-F2. Consistent with these findings, FuFA-F2 prevented the induction of *Myostatin*, *Activin A*, *MAFbx* and *MuRF1*, and attenuated muscle fibrosis and local inflammation.

**Conclusions:** These findings demonstrate that FuFA-F2 preserves skeletal muscle mass and function in the C26 model of cancer cachexia, despite ongoing tumor progression. FuFA-F2 markedly attenuates tumor-induced transcriptional reprogramming and associated catabolic, inflammatory and fibrotic responses, supporting its potential as a therapeutic strategy to preserve skeletal muscle during cancer cachexia.

## 1. INTRODUCTION

Cachexia is a complex, multifactorial syndrome frequently observed in cancer patients, characterized by progressive weight loss, muscle atrophy, and systemic inflammation, which can also be associated with significant adipose tissue atrophy and anorexia. Cachexia is linked to functional decline, diminished quality of life, and decreased tolerance and responsiveness to anticancer treatments [1]. In advanced cancer, the loss of skeletal muscle mass is recognized as an independent predictor of mortality [2]. Thus, cachexia and muscle wasting in particular represent a major challenge for cancer patients care. Unfortunately, to date, there is no effective treatment, although a clinical trial is currently underway with Ponsegromab targeting the anorexigenic GDF15 cytokine [3]. The mechanisms driving cancer cachexia are not yet fully understood, but result from complex interactions between tumor-derived factors [1, 4], systemic inflammation [5], metabolic alterations [6], and a disrupted balance between muscle protein synthesis and degradation [1]. The imbalance between protein breakdown and synthesis, which leads to muscle atrophy, is a central feature of cancer cachexia. Muscle protein degradation, primarily mediated by the ubiquitin-proteasome system, is regulated by two key atrogenes: the muscle-specific E3 ubiquitin ligases MAFbx and MuRF1 [7] which are upregulated during muscle atrophy and cachexia in murine models [8]. Conversely, protein synthesis is regulated by members of the TGF-β superfamily, such as Myostatin (Mstn) [9] and Activin A (ActA) [10], which are potent negative growth factors of muscle mass. Numerous studies emphasize that ActA and Mstn contribute to muscle atrophy induced by some types of cancers [11–13].

Furan fatty acids (FuFAs) are natural compounds produced by plants [14] and microorganisms [15, 16], found in a wide range of foods [17]. Fish represents a particularly rich source of FuFAs, as it consumes algae and marine microorganisms abundant in these fatty acids [18, 19]. FuFAs are also present in butter and dairy products [20, 21], cereals, vegetables, and fruits [18]. Notably, the FuFA-F2 form (10,13-epoxy-11-methyl- octadecan-10,12-dienoic acid), also called 9M5, is especially abundant in the latex of the PB235 clone of *Hevea brasiliensis*, used in natural rubber production [22, 23]. In Germany, the average daily intake of FuFAs from foods such as fish, milk, and soybean oil is estimated to range from 10 to 40 mg [21]. In populations with a high consumption of seafood and fermented dairy products, such as the Mediterranean population, this daily intake is likely to be higher.

FuFAs exhibit a range of biological activities that highlight their potential as bioactive compounds beneficial to human health [24]. Owing to their electron-rich furan ring, FuFAs possess notable antioxidant properties [25–27]. When exposed to reactive oxygen and nitrogen species (RONS), the furan ring opens, forming an unstable and highly reactive dioxoene intermediate [28]. *In vitro*, studies have shown that FuFAs enhance the expression of adiponectin, an adipokine known for its anti-inflammatory and insulin sensitive effects in 3T3-L1 adipocyte cells [29]. *In vivo*, using a rat model of arthritis, FuFAs demonstrated stronger anti-inflammatory effects than eicosapentaenoic acid (EPA) [30]. Moreover, while many studies attribute the cardioprotective effects of fish consumption primarily to polyunsaturated fatty acids (n-3 PUFAs) [31, 32], some evidence suggests that FuFAs may also contribute to these benefits [19].

Recently, using FuFA-F2 (9M5), we demonstrated that this fatty acid enhances protein content in C2C12 myotubes *in vitro* [33]. In addition, oral administration of FuFA-F2 for three weeks increased muscle mass and promoted a more oxidative muscle metabolism in three-month-old C57Bl/6J mice [33]. Furthermore, preventive nutritional supplementation with FuFA-F2 for three months in diet-induced obese (DIO) mice led to increased muscle mass and attenuated the development of metabolic disorders [34]. FuFA-F2 supplementation also improved body composition, increased resting energy expenditure, enhanced insulin sensitivity, and prevented hepatic steatosis [34]. More recently, we showed that four weeks of FuFA-F2 supplementation in already obese mice was sufficient to reverse the development of metabolic dysfunction-associated steatotic liver disease (MASLD), increase skeletal muscle mass, and protect against cartilage degradation induced by a high-fat, high-sucrose diet [35]. Together, these results indicate that FuFA-F2 enhances muscle mass across different metabolic conditions and partially recapitulates the beneficial effects of physical exercise.

The aim of this study was to investigate whether FuFA-F2 supplementation could counteract or delay the onset of muscle wasting, both *in vitro* using a C2C12 myotube model of atrophy and *in vivo* in a well-characterized cancer cachexia model consisting of mice bearing C26 adenocarcinoma cells [36].

## 2. MATERIALS & METHODS

### 2.1 FuFA-F2 purification

Fresh latex from the PB235 *Hevea brasiliensis* clone was collected from a plantation of Chachoengsao Rubber Research Center (CRRC), Chachoengsao, Thailand. Lipid extraction was performed by slowly adding the freshly harvested latex (25 l) dropwise into continuously stirred ethyl acetate (120 l, Univar, Illinois, USA) at a latex-to-solvent ratio of ∼1:5 (v/v) in a 500-l stainless Steel Tank (B.E. Marubichi, Thailand). The coagulum, primarily consisting of insoluble polyisoprene, was removed and the extract was left to decant for 2 hours. The upper layer, consisting of lipid-rich ethyl acetate, was collected, evaporated, and filtered to remove precipitates (Whatman No.1, England). Total lipid extract (∼125 g, yield ∼0.5% w/w) was further purified using flash chromatography (Biotage Selekt, Sweden). For this, a lipid extract (30 g) was loaded onto a 350 g silica gel (100Å 60 µM, Biotage Sfär Silica D – Duo 60 µM). The mobile phase was made of hexane (100% from 0-15 min) followed by a gradient of hexane:diethyl ether (90:10, v/v) for 5 min (from 15-20 min) with a final 30 min plateau at the same ratio (from 20-50 min), all at a flow rate of 50 ml/min and with a collection volume of 50 ml/fraction. The quality of the separation was assessed by spotting aliquots of each fraction onto a thin layer chromatography plate, following the method described by Liengprayoon et al. for neutral lipids [37]. The fractions with the highest trifuranoylglycerol (TFG) purity were analyzed by GC-FID after derivatization to methyl esters, as previously described [37]. The FuFA-F2 methyl ester purity within all obtained methyl esters, calculated from the GC-FID, was approximately 98%.

### 2.2 Cell Culture and Treatments

Mouse myoblasts of the C2C12 cell line (ATCC) were seeded at a plating density of 7000 cells/cm^2^ in 6- well plates. They were grown in DMEM high glucose (4.5 g/l) supplemented with gentamycin (50 µg/ml), amphotericin (50 µg/ml), and fetal calf serum (10%). Terminal differentiation was induced at cell confluence by lowering the medium serum concentration (2%). On day 5, a cocktail composed of TNFα and INFγ at 10 ng/ml was added to the culture medium to induce myotube atrophy. When indicated, FuFA-F2 was added to the culture medium at a final concentration of 10 µM.

### 2.3 Cytoimmunofluorescence

Cytoimmunofluorescence myoblast differentiation was assessed by observation of morphological changes and accumulation of muscle-specific markers. After methanol fixation and three washes with PBS-gelatin (0.2%), cells were labeled with an antibody raised against Troponin T (T6277, Sigma, diluted at 1:50). Nuclei were stained with Hoechst 33258 (1 µg/ml). Myotube surface area analysis was performed automatically using Fiji software by counting troponin T-positive cells to obtain the total myotube surface area per image.

### 2.4 Cancer cachexia model in vivo

3-month-old male CD2F1 mice were randomly divided into three groups using variables weight, lean and fat masses (Randomice v1.1.7) [38]: control group without tumor (Control group, 6 mice), C26 tumor-bearing mice group (C26 group, 10 mice), and mice carrying C26 tumors that were supplemented with an average of 13 mg/kg of FuFA-F2 (C26 + FuFA group, 10 mice). 500,000 C26 tumor cells were subcutaneously injected in the fat pad of the mouse as previously described [36]. The mice were housed in pairs in adapted TIII cages (Tecniplast, Italy) with a running wheel (23-cm diameter, Intellibio) for each animal as previously described [39]. They were maintained on a 12-hour light/dark cycle (lights on at 7:30 am). Food (A04, Safe, Augy France) and water were provided ad libitum. Enrichment (wood shavings and polycarbonate red tunnel) was provided in each cage. Mice were treated once daily by oral gavage with vehicle (50% PEG-400, 0.5% Tween 80%, and 49.5% distilled water) or FuFA-F2 (20 mg/kg) as described in Figure 2B. Mice body weight and food consumption were determined every day. Our institution’s guidelines for the care and use of laboratory animals were followed and all experimental procedures were approved by the local ethical committee in Montpellier (CEEA-LR 36) and French ministry of higher education, research and space, France (Reference APAFIS #47638-2024022013115190).

### 2.5 Spontaneous wheel running

The spontaneous and voluntary activity of the mice was measured using an activity wheel of 23 cm of diameter (ActiviWheel, Intellibio) for each animal. Running distances were recorded continuously throughout the 14 days of the experiment using the ActiviWheel software v5.1 (Intellibio).

### 2.6 Body composition

Mice’s whole-body composition (fat and lean masses) was measured as indicated in Figure 2B using the minispec Live Mice Analyzer (LF90 Bruker), according to the manufacturer’s instructions.

### 2.7 Histological study

Tibialis anterior muscles were embedded in a cryoprotective medium and rapidly frozen in isopentane cooled with dry ice (T < −80 °C). Samples were stored at −80 °C until use. Cryosections (10 µm thickness) were prepared and used for fluorescent labeling of muscle fiber membranes with Wheat Germ Agglutinin conjugated to Alexa Fluor 488 (WGA). Sections were rehydrated in 1× phosphate-buffered saline (PBS) and incubated with WGA (6 µg/mL) for 10 min at 37 °C. After washing in PBS, slides were mounted with Permafluor mounting medium and scanned using an Axio Scan slide scanner (Zeiss, MRI, INM). Muscle fiber segmentation was performed using the Cellpose algorithm, a deep-learning–based segmentation algorithm. Image J free software was used to analyze and quantify the pictures for each entire area.

For fibrosis analysis, sections were fixed in 4 % paraformaldehyde for 10 min and rinsed in 1× PBS. Slides were then incubated for 1 h at room temperature in a 0.01 % Sirius Red solution (Sirius Red F3B, CI 35780, Sigma) prepared in 1.3 % picric acid (Analytic Lab). After staining, sections were washed in 0.5 % acidified water containing glacial acetic acid and allowed to dry completely. Slides were then dehydrated through two baths of 100 % ethanol followed by two baths of xylene and mounted with coverslips using Eukitt mounting medium. Whole-slide images were acquired using a Hamamatsu slide scanner (MRI, INM). Fibrotic areas were quantified using NDP.view2, a free slide-viewing and analysis software.

### 2.8 Mitochondrial enzymatic activities

Citrate synthase and cytochrome c oxidase (Complex IV) activities were measured in quadriceps muscle as previously reported [40, 41].

### 2.9 Measurement of inflammatory cytokines in plasma

At the end of the experiment, the animals were anaesthetized and blood was collected into heparinized tubes. Plasma was collected and stored at –80 °C until analysis. Cytokine plasma levels were detected using the bead-based LEGENDplexTM Mouse Cytokine Release Syndrome Panel (13-plex) immunoassay from BioLegend according to the supplier’s suggestions. Data was acquired using the CytoFLEX (Beckman Coulter) flow cytometer.

### 2.10 Quantitative Analysis of Furan Fatty Acid (FuFA-F2)

Muscle, liver, and feed samples were dried in the freeze dryer (Lyovac GT2R, Leybold) for 4-5 days. Feed samples (each ∼5 g) were extracted twice with 50 ml *n*-hexane for 10 min in the ultrasonic bath (Sonorex, Super RK 106, Bandelin). The extracts were combined in a 100 ml pear-shaped flask, and the solvent was removed by using a rotary evaporator (300 mbar, 38 °C, Laborata 4002 control, Heidolph). The residue was transferred with *n*-hexane in a 4 ml brown glass vial and the solvent was evaporated under a gentle stream of nitrogen at 38 °C (EC 1, Köhler-VLM). After the residue was weighed (∼300 mg), the vial was filled up to 3 mL with *n*-hexane (∼100 mg fat/ml). Transesterification of all samples was done according to Bauer *et al*. with slight changes [42]. For muscle and liver samples, ∼15 mg (corresponding to 1-2 mg fat) of the dried samples were weighed into a brown glass tube, while for plasma samples an aliquot of 100 µl plasma was used. To each sample, the internal recovery standard (25 µl of M-11-5, 80 µg/mL in *n*-hexane) as well as 3 ml of freshly prepared 1% sulfuric acid in MeOH were added. With argon (5.0) used as a protection gas, the samples were transesterified at 80 °C for 2.5 h. After that, the tube was cooled on ice, 1 ml of demineralized water and 1 ml of saturated sodium chloride solution were added, and the generated methyl esters (ME) were extracted with 1 ml of *n*-hexane. The phase separation was improved using a centrifuge (1200 rpm, 2 min). For the determination of the FAME content (except for plasma samples), 0.5 ml of the hexane phase was transferred into a 1.5 ml brown glass vial, the solvent was evaporated, the residue was weighed, and the FAMEs were again diluted in 0.5 ml of *n*-hexane. An aliquot of 75 µl of the sample solutions (500 µl in case of plasma samples) was diluted in a total volume of 1000 µl with hexane while adding 25 µl of a mix of the internal quantification standard (9M5-ethyl ester (EE), 3.06 µg/ml in hexane) and the syringe standards (14:0- EE and 22:0-EE, both ∼0.01 mg/ml in hexane). Samples were measured against a matrix standard solution, which was prepared the same way, except that the internal recovery standard was added in the last dilution step, resulting in a final concentration of 0.14 µg/ml M-11-5.

For transesterification of the feed extract, an aliquot of 30 µl (corresponding to ∼3 mg fat) was taken and filled into a brown glass tube. After adding the internal recovery standard (50 µl of M-11-5, 80 µg/ml in hexane), the transesterification was done as above, except for an ME-extraction with 2 ml hexane and a dilution of the sample solution by a factor of 40.

Generated FuFA-ME were measured on a 6890 Series II/5973 MSD/6890 ALS GC-MS system (Hewlett-Packard/Agilent). GC parameters were set according to the method of Wendlinger *et al*. [43] except for a transfer line temperature of 270 °C, and ions and time windows measured in the selected ion monitoring (SIM) mode were adapted from Bauer *et al*. [42]. Determination of the recovery (calculated with M-11-5) and internal quantification of 9M5-ME (calculated with 9M5-EE for muscle samples and M-11-5 for liver and plasma samples) were done at *m/z* 165.1. Detection limit and limit for quantification of 9M5-ME were 5.15 pg and 17 pg, respectively.

### 2.11 Protein isolation and western blotting analysis

Frozen gastrocnemius muscle samples were homogenized using an Ultra Turax homogenizer. Protein levels were assessed by Western blotting. Total proteins were measured using the Bio-Rad protein assay according to the manufacturer’s instructions. 40 µg of each protein extract were loaded on stain-free 4%–20% precast gels (Bio-Rad) for protein separation by electrophoresis, followed by transfer to nitrocellulose membranes (Trans-Blot Turbo Blotting System; Bio-Rad). Signals were revealed using a ClarityTM Western ECL Substrate Kit, and proteins were visualized by enhanced chemiluminescence using the ChemiDoc Touch Imaging System and quantified with Image Lab. Touch Software (version 5.2.1). The Stain-Free technology was used as a loading control.

Primary antibodies used include: anti-Phospho-AMPKα (Thr172) (1/1000, Cell Signaling #2535); anti- AMPKα (1/1000, Cell Signaling #2532); anti-LC3A (1/1000, Cell Signaling #4599); anti-SQSTM1/p62 (1/1000, Cell Signaling #5114). Library preparation is performed using Optimal Dual-mode mRNA Library Prep Kit

### 2.12 RNA-seq analysis

#### RNA extraction, library preparation and sequencing

Total RNA was extracted from gastrocnemius muscle samples using TRIzol® reagent, and cDNA was synthesized using the PrimeScript 1st Strand cDNA Synthesis Kit (Takara Bio). Double-stranded cDNA libraries were generated using dUTP in place of dTTP during second-strand synthesis. The resulting cDNA fragments were subjected to end repair and 3′ A-tailing, followed by ligation of sequencing adapters. The ligated products were amplified by PCR and subjected to quality control. Following denaturation, single-stranded library molecules were circularized, and residual linear DNA was digested. The resulting single-stranded circular DNA molecules were amplified by phi29-mediated rolling circle amplification (RCA) to generate DNA nanoballs (DNBs), each containing multiple copies of the original library molecule. DNBs were loaded onto a patterned nanoarray, and paired-end sequencing (PE150) was performed on a G400 platform (BGI-Hong-Kong, China).

#### RNA-seq data processing and analysis

Sequencing data were filtered using SOAPnuke [44] to remove reads containing sequencing adapters, reads with more than 20% low-quality bases (base quality ≤15), and reads with more than 5% undetermined bases (N). The resulting clean reads were stored in FASTQ format and subsequently analyzed using the Dr. Tom Multi-omics Data Mining System (BGI). Clean reads were aligned to the reference genome using HISAT2 [45]. Fusion genes and differential splicing events were identified using Ericscript v0.5.5 [46] and rMATS v4.1.2 [47], respectively. For gene-level quantification, clean reads were aligned to a comprehensive gene set, including known and novel coding and non-coding transcripts, using Bowtie2 [48]. Gene expression levels were quantified using RSEM v1.3.1 [49]. Differential gene expression analysis was performed using DESeq2 v1.34.0 [50]. Differentially expressed genes (DEGs) were identified using a false discovery rate (FDR) or Q-value threshold of 0.05. Heatmaps were generated using the pheatmap package v1.0.12. Gene Ontology (GO) and Kyoto Encyclopedia of Genes and Genomes (KEGG) enrichment analyses were performed using Phyper based on a hypergeometric test. Enrichment significance was assessed after multiple-testing correction, with Q ≤ 0.05 considered statistically significant (http://github.com/jdstorey/qvalue).

### 2.12 Gene Expression Studies

Total RNAs were extracted from gastrocnemius muscle samples using Trizol, and cDNAs were generated using the PrimeScript-1st strand cDNA Synthesis Kit (Takara Bio). Real-time PCR was performed using Owi Green FAST qPCR Premix (OWIBIO) and LightCycler 480 (Roche). Gene expression was normalized to the expression of the housekeeping gene Rpl13. Primer sequences are listed in Supplemental Figure S1.

### 2.12 Statistical analyses

All results are presented as means ± SD, or as percentages. Groups were tested by a one-way ANOVA test. The limit of statistical significance was set at p<0.05. The means with different letters are significantly different. Statistical analyses were performed using the StatView software.

## 3. RESULTS

### 3.1 FuFA-F2 prevents muscle atrophy of C2C12 myotubes induced by TNFα and IFNγ

First, we evaluated the effects of FuFA-F2 *in vitro* on a model of differentiated C2C12 myotubes atrophy in response to combined IFNγ and TNFα treatment [51, 52]. Following myotube differentiation, exposure to IFNγ and TNFα promoted C2C12 myotubes atrophy, as demonstrated by immunofluorescence analysis and quantification of myotubes area (Figure 1A-B). Concomitant treatment with FuFA-F2 prevented cytokine-induced muscle wasting (Figure 1A-B) and slightly increased the myoblast fusion index (Figure 1B).

**Figure 1.**
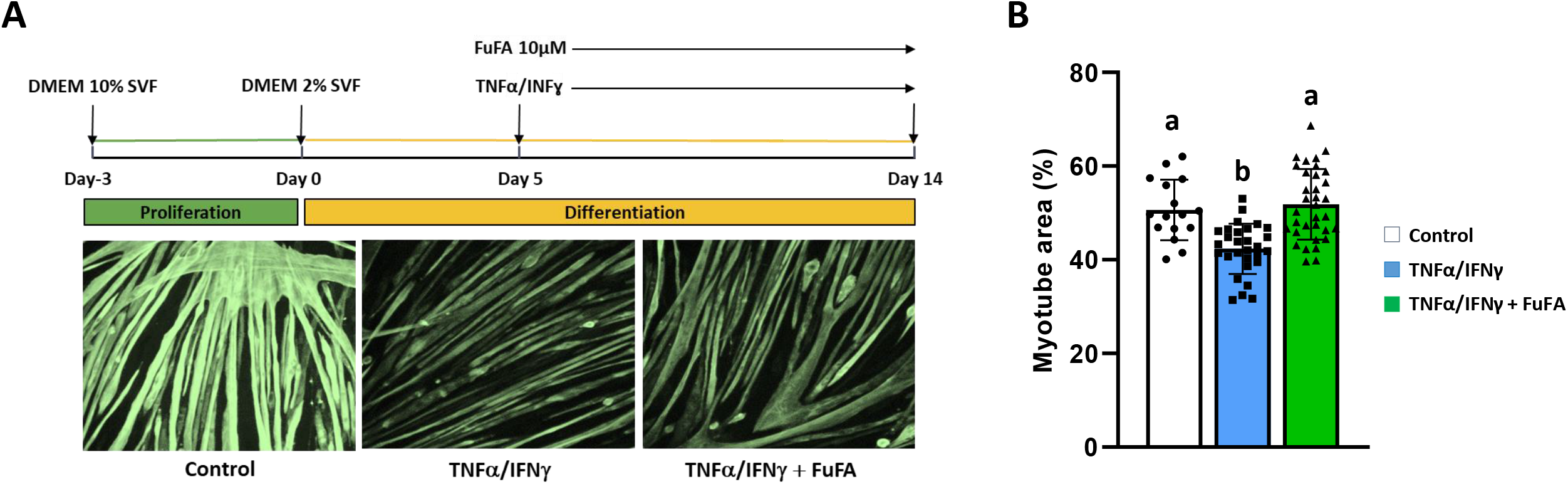
FuFA-F2 prevents muscle wasting of C2C12 myotubes induced by TNFα and IFNγ. (A) Scheme of the experiment and representative pictures of C2C12 myotubes after 14 days of differentiation (Cytoimmunofluorescence studies using an antibody raised against Troponin T). (B) Quantification of myotube area after troponin T staining (%). Results were expressed as means ± SD. Groups were tested by a one-way ANOVA test. The limit of statistical significance was set at p<0.05. The means with different letters were significantly different.

**Figure 2.**
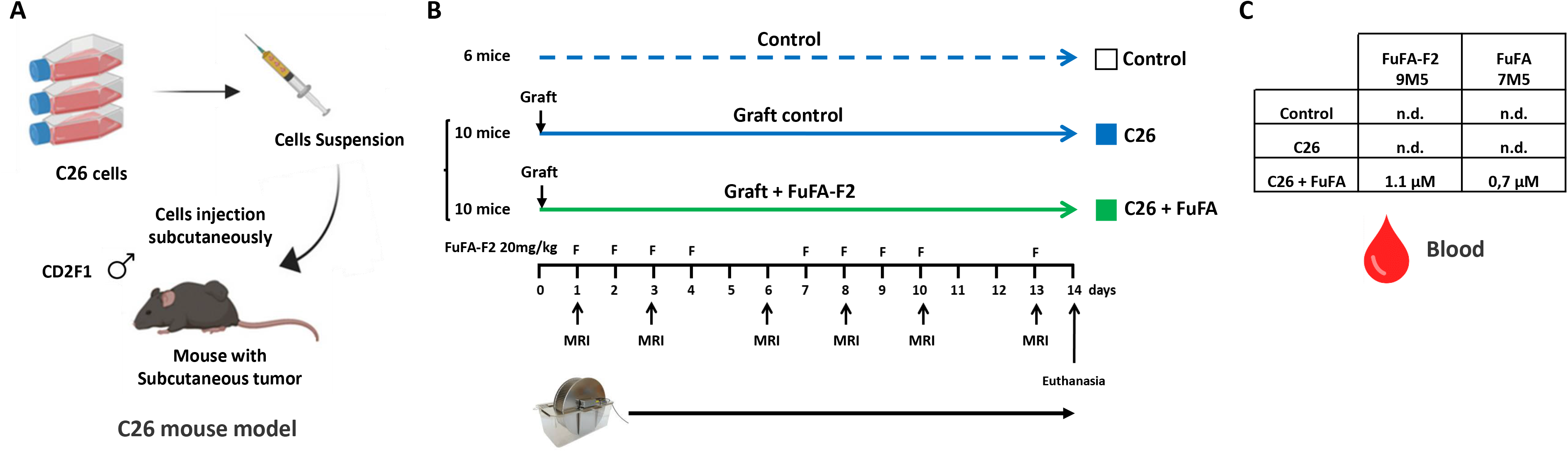
Design of the study in C26 tumor-bearing mice. (A) Scheme of the obtention of C26 tumor-bearing mice. (B) Design of the study. Our experiments were performed on three groups: control group without tumor (Control group, 6 mice), C26 tumor-bearing mice group (C26 group, 10 mice), and C26 tumor-bearing mice supplemented with FuFA-F2 (C26 + FuFA group, 10 mice). When indicated with the label F, mice received by oral gavage the vehicle or FuFAF2 (20 mg/kg). Whole-body composition was measured as indicated by the label MRI using a minispec Live Mice Analyzer (LF90 Bruker). Spontaneous physical activity was recorded throughout the study using in-cage running wheels. (C) Concentration of FuFA-F2 (9M5) and its derivative FuFA 7M5 in the blood of animals after 14 days of treatment. n.d. not detected.

### 3.2 FuFA-F2 improved cancer cachexia symptoms of C26 tumor-bearing mice

To confirm the *in vitro* results obtained in C2C12 myotubes, the effects of FuFA-F2 were next evaluated in mice bearing C26 tumors, a well-established model of cancer cachexia [36]. Starting one day after C26 cells transplantation, mice received daily oral gavage of either vehicle or FuFA-F2 (20 mg/kg) administered every day except weekends for two weeks, corresponding to an average dose of 13 mg/kg/day (Figure 2A–B). To verify the effective absorption of FuFA-F2 (9M5), plasma concentrations of FuFA-F2 and one of its degradation products, FuFA 7M5 (7-(3-methyl-5-pentylfuran-2-yl)-heptanoic acid) [29], were measured. As expected, after two weeks of supplementation, plasma levels reached 1.1 µM for FuFA-F2 and 0.7 µM for FuFA 7M5 in treated mice, whereas both compounds were undetectable in control and untreated C26 mice (Figure 2C). FuFA-F2 treatment attenuated body weight loss without altering tumor growth (Figure 3A-C). We observed that FuFA-F2 administration significantly reduced tumor-free body weight loss (Figure 3D-E) Importantly, cumulative food intake did not differ significantly between the groups (Figure 3F).

**Figure 3.**
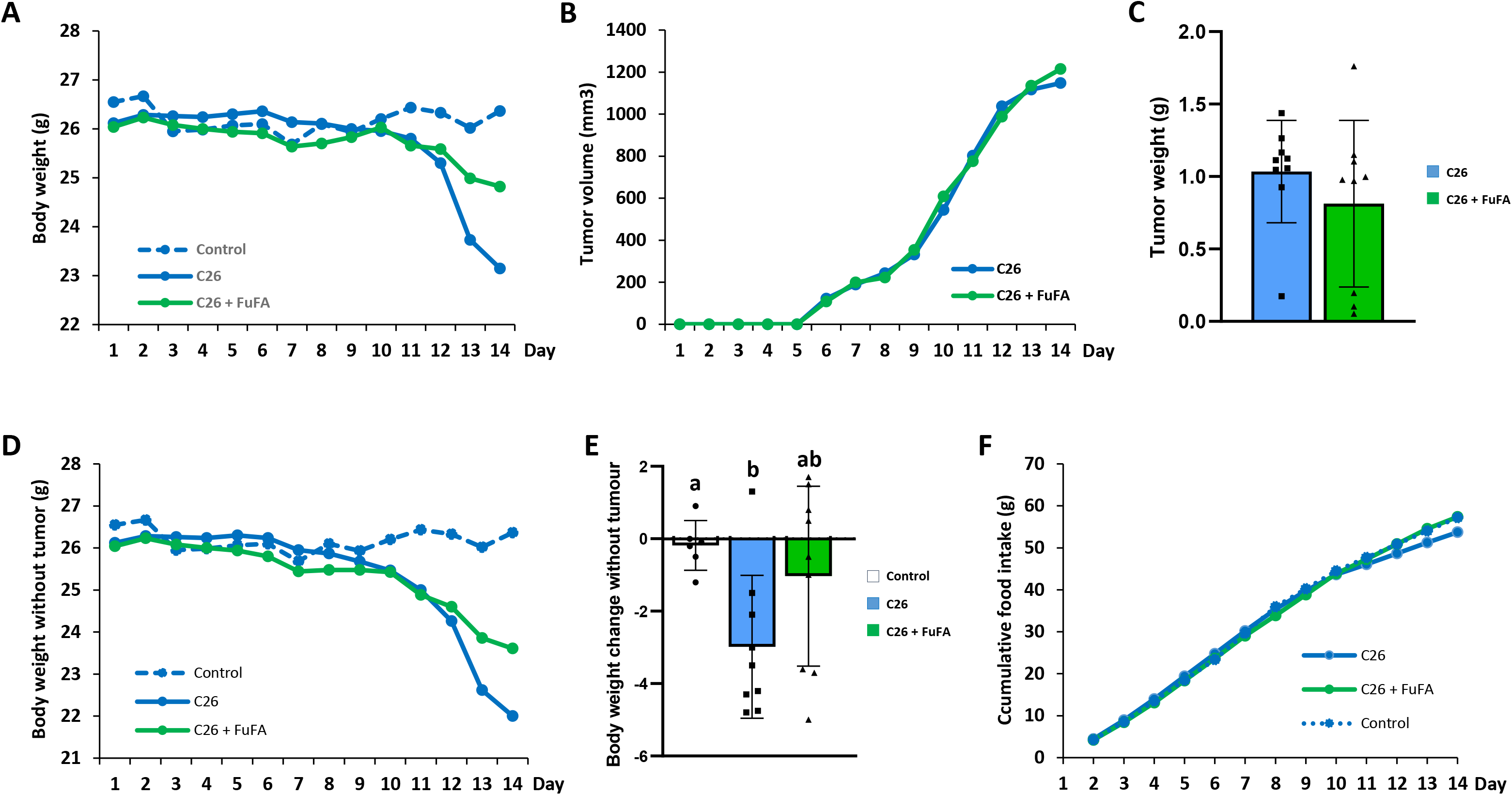
FuFA-F2 improved cancer cachexia symptoms of C26 tumor-bearing mice. (A) Evolution of body weight with tumor in the different groups of mice. (B-C) Tumor volume (B) and tumor weight (C) of mice after euthanasia. (D-E) Body weight without tumor is shown across time (D) or as total change at the point of euthanasia (E). (F) Accumulative food intake of mice. Results were expressed as means ± SD. Groups were tested by a one-way ANOVA test. The limit of statistical significance was set at p<0.05. The means with different letters were significantly different.

### 3.3 FuFA-F2 prevented muscle wasting but did not inhibit fat loss of C26 tumor-bearing mice

Body composition measurements taken throughout the study (on days 1, 3, 6, 8, 10, and 13) revealed that FuFA-F2 supplementation did not prevent the substantial loss of fat tissue in C26 tumor-bearing mice (Figure 4A-B). At euthanasia on day 14, both treated and untreated C26 model groups showed almost complete disappearance of the epididymal white adipose tissue (eWAT). To further evaluate the well-being and quality of life of the mice, we monitored not only standard parameters such as body weight and prostration but also spontaneous physical activity using in-cage running wheels. During the first 10 days, no significant differences in the distance traveled were observed between the groups (Figure 4C). However, over the final four days, mice in the C26 model group exhibited a sharp decline in activity compared to non-grafted controls (Figure 4D). Notably, C26 tumor-bearing mice supplemented with FuFA-F2 maintained their physical activity levels (Figure 4D), suggesting preserved functional capacity and overall well-being.

**Figure 4.**
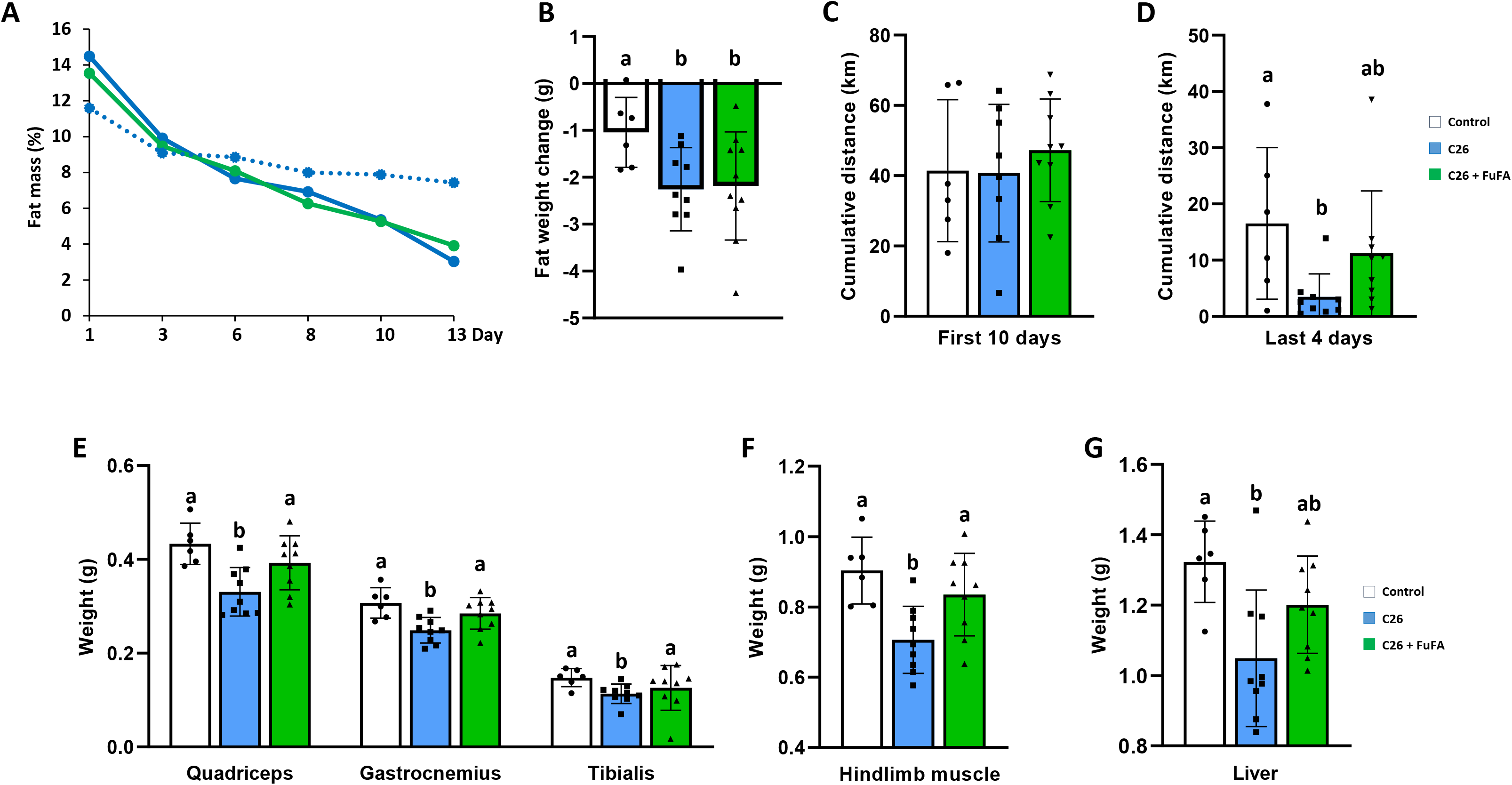
FuFA-F2 prevented muscle wasting but did not inhibit fat loss of C26 tumor-bearing mice. (A-B) Fat mass evolution measured using a minispec Live Mice Analyzer (LF90 Bruker) expressed as % of total mass (A) or shown as weight change (g) at the point of euthanasia. (C-D) Cumulative distance (km) covered during the first 10 days (C) or the last 4 days (D). (E-G) Organ weight expressed in g for the quadriceps, gastrocnemius, and tibialis muscles (E), hindlimb muscle (F), and liver (G). Results were expressed as means ± SD. Groups were tested by a one-way ANOVA test. The limit of statistical significance was set at p<0.05. The means with different letters were significantly different.

Following euthanasia on day 14, and consistent with the pronounced sarcopenia in this cachexia model, the weights of quadriceps, gastrocnemius, and hindlimbs muscles were markedly reduced in the C26 model group (0.706 g ± 0.095 g) compared to non-grafted controls (0.903 g ± 0.095 g) (Figure 4E-F). In contrast, C26 tumor-bearing mice supplemented with FuFA-F2 exhibited muscle weights (0.835 ± 0.117 g) that were no longer significantly different from those of non-grafted controls (0.903 ± 0.095 g) (Figure 4E-F). Interestingly, another key metabolic organ, the liver, also appeared to be protected by FuFA-F2 supplementation (Figure 4G). Liver weight was markedly reduced in C26 mice (1.049 g ± 0.065 g) compared with controls (1.323 g ± 0.047 g), whereas this decrease was attenuated in the C26 mice receiving FuFA-F2 (1.201 g ± 0.046 g).

### 3.4 FuFA-F2 prevented muscle wasting but only partially counteracted muscle remodeling of C26 tumor-bearing mice

To determine whether the preservation of muscle mass induced by FuFA-F2 was associated with changes in the metabolic profile of muscle fibers, we measured mitochondrial activity in the quadriceps. Citrate synthase (CS) activity, commonly used as a marker of mitochondrial content, as well as the activity of mitochondrial respiratory chain complex IV, remained unchanged across all groups (Figure 5A-C).

**Figure 5.**
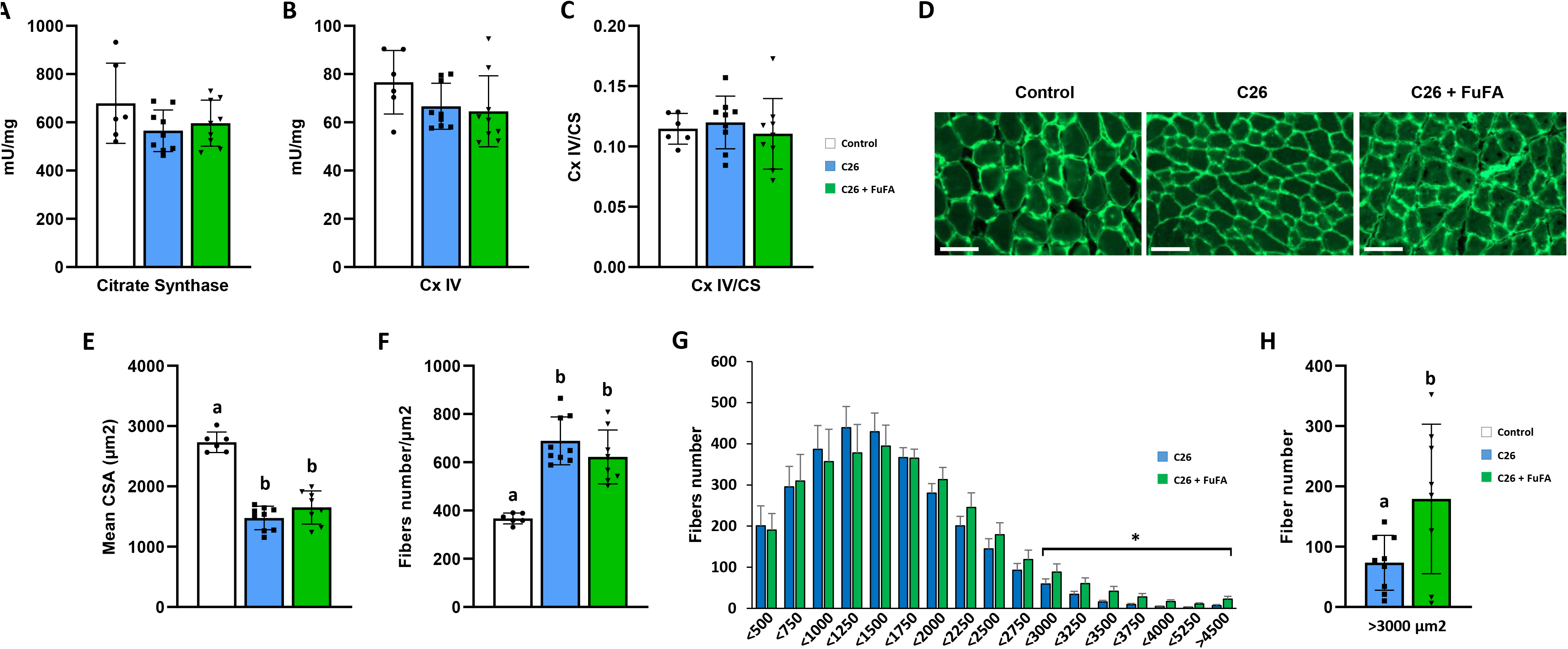
Mitochondrial activity and CSA analysis. (A-C) Mitochondrial activity from quadriceps muscle extracts in the different experimental groups: (A) Citrate synthase activity; (B) Mitochondrial respiratory chain activity of complex IV; (C) Mitochondrial respiratory chain activity of complex IV reported to Citrate Synthase activity (CS). (D) Representative pictures of transverse cross-sectional area of tibialis muscle after WGA staining. Scale bar, 100 µm. (E-H) Quantifications from images shown in D and evaluating the mean fibers area expressed in µm^2^ (E), total fibers number by µm^2^ (F), fiber size distribution in tibialis muscles (G), and the number of fibers with CSA greater than 3000 µm^2^ (H). Results were expressed as means ± SD. Groups were tested by a one-way ANOVA test. The limit of statistical significance was set at p<0.05. The means with different letters were significantly different.

Next, cross-sections of the tibialis muscle were analyzed (Figure 5D). Morphometric analysis revealed a pronounced reduction in the mean cross-sectional area (CSA) of muscle fibers in mice bearing C26 carcinoma, regardless of FuFA-F2 treatment (Figure 5E). Conversely, the apparent number of muscle fibers was increased (Figure 5F). However, detailed analysis of fiber size distribution showed that FuFA-F2 supplementation partially preserved the population of larger muscle fibers (Figure 5G). Specifically, FuFA- F2 treatment led to a significant increase in fibers exceeding 3000 µm² (+244%, *p* < 0.05) compared with the C26 group (Figure 5H). These findings indicate that FuFA-F2 supplementation partially counteracts the muscle remodeling induced by C26 carcinoma.

### 3.5 FuFA-F2 attenuates tumor-induced transcriptional reprogramming of skeletal muscle

To better understand the role of FuFA-F2 in preserving muscle mass despite tumor growth, we performed RNA-seq analysis on RNA extracted from the quadriceps muscles (Figure 6A). Transcriptomic analysis revealed profound reprogramming of muscle tissue in response to tumor growth. Comparison of the Control and C26 groups identified 5,465 differentially expressed genes (DEGs) out of 17,069 detected genes (32%), indicating extensive disruption of the muscle transcriptional program induced by tumor growth (Figure 6C and 6E). Remarkably, this transcriptional response was markedly attenuated by FuFA-F2, with only 366 DEGs (2%) detected between Control and C26+FuFA-F2 mice (Figure 6F and 6H), indicating that FuFA-F2 largely preserved the transcriptional profile of control muscle. This observation was further supported by principal component analysis (PCA), which showed a clear separation between Control and C26 groups, whereas C26+FuFA-F2 samples clustered closer to Control samples (Figure 6B). Closer examination of the KEGG pathways altered by tumor growth (Control vs. C26) (Figure 6D) revealed enrichment in numerous pathways, including those related to protein synthesis (Ribosome biogenesis, ribosomes), catabolism and protein degradation (proteasome, autophagy, mitophagy), signaling pathways controlling core metabolism (AMPK, FoxO, and mTOR signaling pathways), and signaling pathways previously implicated in muscle atrophy during cachexia such as the TGF-beta pathway (the atrophic role of Activin-A and Myostatin), and the proinflammatory NF-Kappa beta pathway (Supplemental figure 2-11). Some of these transcriptomic signatures could represent adaptative responses, for instance with genes coding for ribosomal proteins and translation initiation factors which were up-regulated in the catabolic muscles of C26 mice. They also suggest that the competence and sensitivity of cachectic muscles to various signaling pathways were profoundly affected with increased expression of different genes encoding key signaling pathways proteins such as Smads, mTor, Akt, Foxo, Myd88, or Rel proteins, even though the actual signaling output (activation or inhibition in C26 muscles) remained complicated to predict. The muscle bulk RNA-seq profile also revealed the up-regulation in C26 mice of genes specifically expressed by immune cells (macrophages, neutrophils…), such as Clcf1, Ly6c1, Lcn2, Cxcl13 and Ccl8, suggesting increased immune cell presence in the tissue, consistent with previous reports [53–55]. Strikingly, these pathways were largely preserved in the Control versus C26+FuFA-F2 comparison (Figure 6G), with only minimal transcriptional modulation, supporting the ability of FuFA-F2 to preserve a control-like muscle transcriptomic profile in tumor-bearing mice. We next compared the C26 and C26+FuFA-F2 groups and identified 3,932 DEGs, corresponding to 23% of the detected genes (Figure 6I and 6K). Notably, 3,428 of these genes were also differentially expressed between Control and C26, representing approximately 87% of the genes differentially expressed between C26 and C26+FuFA-F2. This substantial overlap indicates that the transcriptional changes associated with FuFA-F2 treatment predominantly involve genes affected by tumor growth, and relatively upstream in the cascade of events that will lead to muscle atrophy. Consistent with this finding, the KEGG pathways altered in the Control versus C26 and C26 versus C26+FuFA-F2 comparisons (Figure 6J) showed a highly similar pattern (Supplemental figure 14-25). Furthermore, many immune cell markers observed in C26 muscles were also absent after FuFA-F2 treatment suggesting smaller immune infiltrate. Together with the PCA showing that C26+FuFA-F2 samples displayed a transcriptomic profile closer to that of Control samples (Figure 6J), these findings support the ability of FuFA-F2 to preserve the muscle transcriptomic state in tumor-bearing mice.

**Figure 6.**
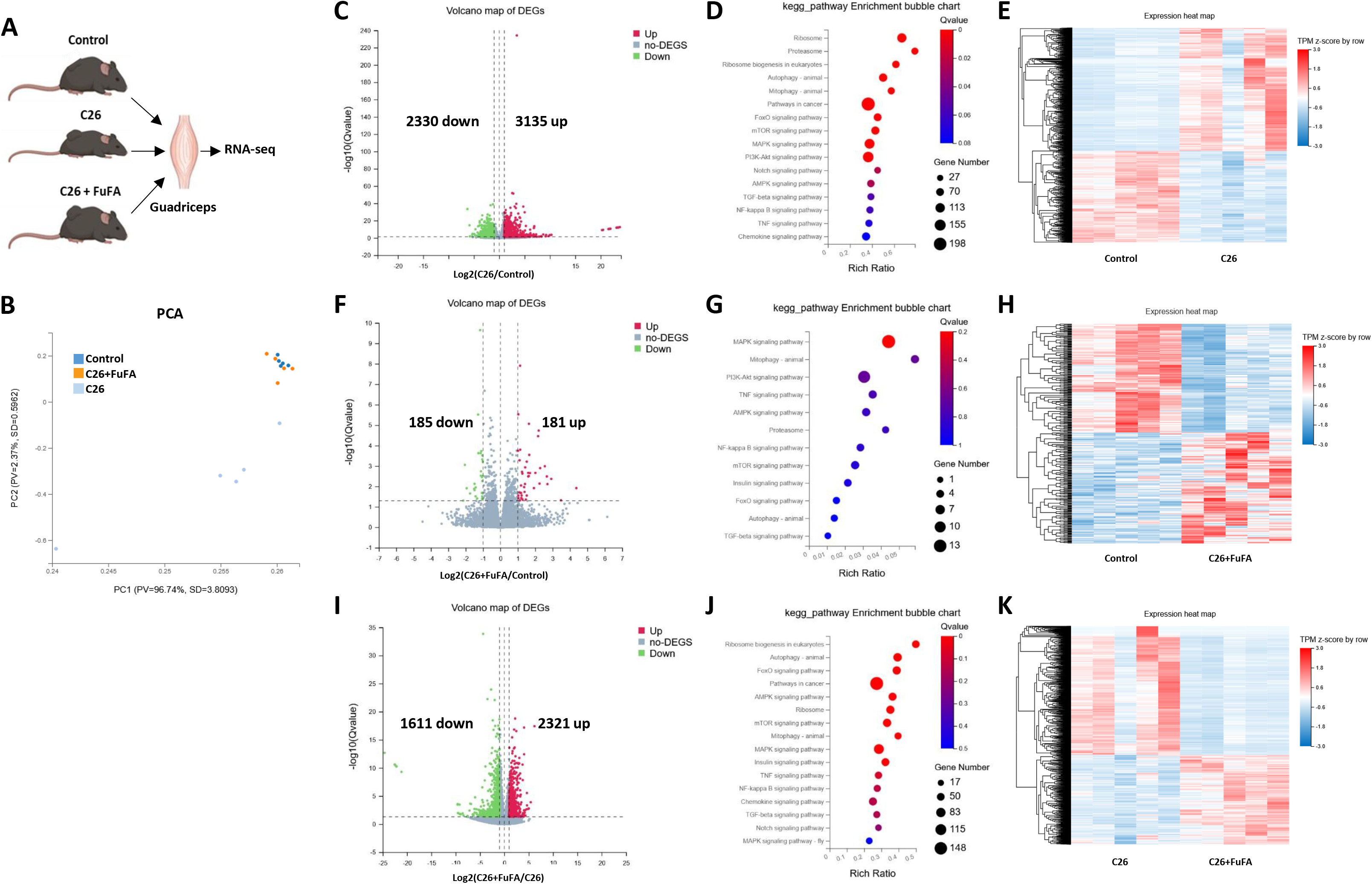
Transcriptomic analysis in skeletal muscle. (A) The transcriptomes of muscle from control, C26, and C26+FuFA-F2 were compared by RNA sequencing (RNA-Seq) (n=5 per group). (B) Principal component analysis (PCA) plot for RNA-seq data of biological replicates of Control, C26, and C26+FuFA. (C) Volcano plot depicting differentially expressed genes in Control vs C26 groups. Red dots represent genes significantly overexpressed in C26 while green dots represent genes underexpressed. (D) KEGG pathway enrichment analysis between Control vs C26. (E) Heatmap Control vs C26. (F) Volcano plot depicting differentially expressed genes in Control vs C26+FuFA groups. Red dots represent genes significatively overexpressed in C26+FuFA while green dots represent genes underexpressed. (G) KEGG pathway enrichment analysis between Control vs C26+FuFA. (H) Heatmap Control vs C26+FuFA. (I) Volcano plot depicting differentially expressed genes in C26 vs C26+FuFA groups. Red dots represent genes significatively overexpressed in C26+FuFA while green dots represent genes underexpressed. (J) KEGG pathway enrichment analysis between C26 vs C26+FuFA. (K) Heatmap C26 vs C26+FuFA.

### 3.6 FuFA-F2 preserves muscle homeostasis during cancer cachexia

Our RNA-seq analysis revealed a profound remodeling of skeletal muscle in C26 mice, with marked alterations in pathways involved in muscle proteostasis including the proteasome (Psmb5, Psmb6, Psmb7…), autophagy (such as elevated levels of Ulk1, Gabarap, Gabarapl1, Atg7, LC3 (Map1lc3b) and p62 (Sqstm1)), mitophagy (Bnip3, Bnip3l (Nix) and Tfeb) and AMPK signaling pathway (Prkaa2, Prkab1 and Prkg3). These pathways are closely associated with the regulation of muscle protein turnover and are known to contribute to muscle wasting during cancer cachexia. We therefore investigated key molecular mediators of these pathways in gastrocnemius muscle by Western blot and qPCR. We first assessed AMPK activation, a key metabolic sensor involved in the regulation of muscle protein turnover and implicated in cancer cachexia [56]. The ratio of phosphorylated to total AMPK was increased in C26 mice compared with controls, indicating enhanced AMPK activation. Notably, this activation was prevented in C26 mice supplemented with FuFA-F2 (Figure 7A–B), suggesting that FuFA-F2 may limit the metabolic alterations associated with muscle wasting.

**Figure 7.**
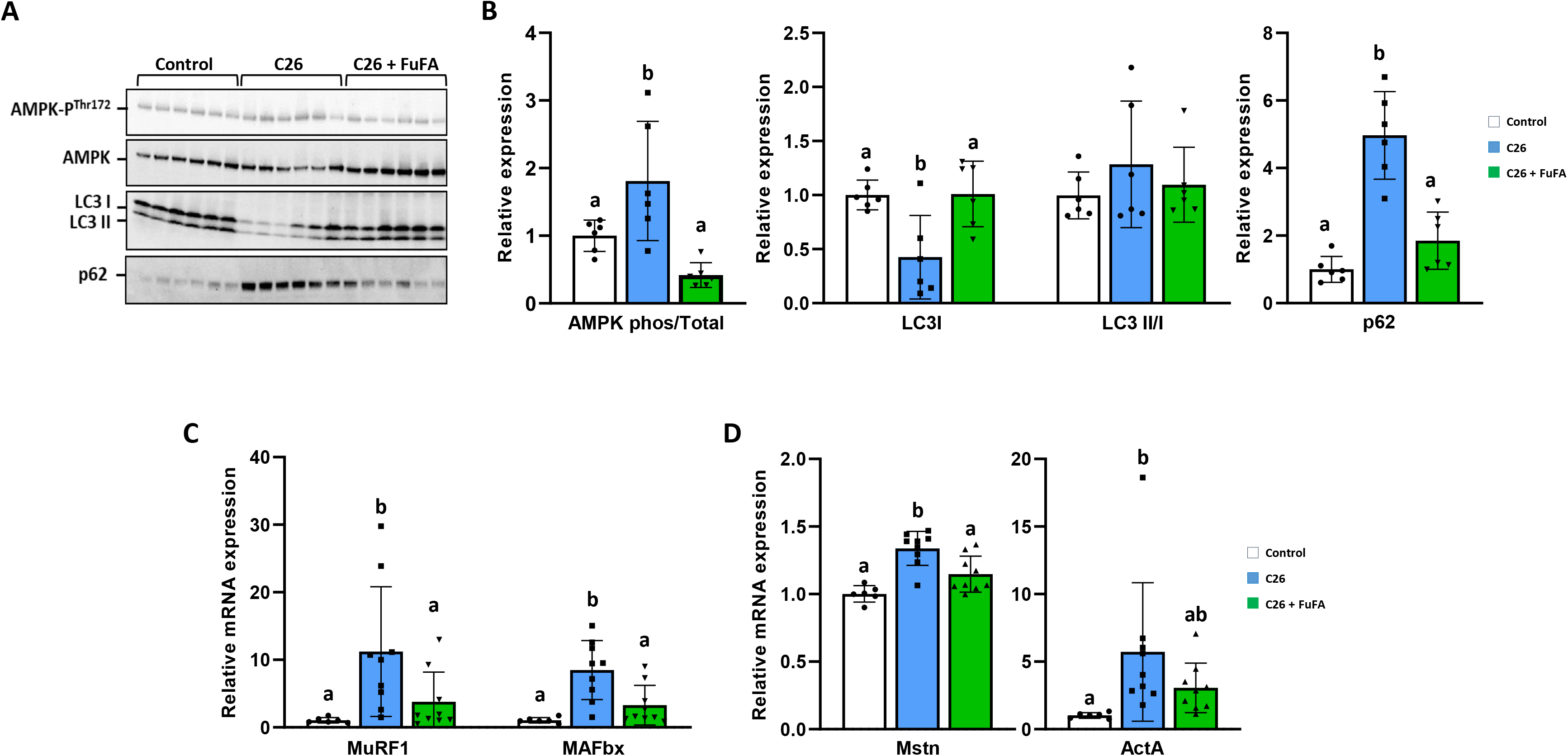
FuFA-F2 prevented muscle loss by suppressing the upregulation of *Myostatin and Activin A*, and the ubiquitin E3 ligases *MAFbx* and *MuRF1*. (A-B) Western-blot analysis of the indicated proteins in gastrocnemius muscle (A, *n* = 6 for each group,) and quantification (B). The Stain-Free technology was used as a loading control. (C-D) Relative mRNA expression in gastrocnemius muscle of *MuRF1* and *MAFbx* (C), or *Mstn* and *ActA* (D). Results were expressed as means ± SD. Groups were tested by a one-way ANOVA test. The limit of statistical significance was set at p<0.05. The means with different letters were significantly different.

We next examined autophagy-related proteins, LC3 and p62, commonly used markers of autophagy [57]. LC3 levels were decreased, whereas p62 levels were increased in C26 muscle compared with controls, and both alterations were prevented by FuFA-F2 (Figure 7A–B). Although previous studies have reported increased autophagy-related markers in C26 muscle [58], the reduced LC3 levels observed here may reflect the advanced stage of cachexia at day 14. However, since LC3 and p62 abundance alone cannot directly assess autophagic flux, this interpretation remains speculative.

Because proteasome-related pathways were also enriched in C26 muscle, we next examined the muscle-specific E3 ubiquitin ligases *MAFbx* and *MuRF1*, key mediators of ubiquitin-dependent protein degradation during muscle wasting [59, 60]. Both genes were strongly upregulated in C26 muscle, whereas their expression remained at control levels in C26 mice supplemented with FuFA-F2 (Figure 7C), consistent with our RNA-seq findings. Finally, we examined *Mstn* and *Acta*, two negative regulators of skeletal muscle mass belonging to the TGF-β superfamily [9, 10]. Both genes involved were upregulated in C26 muscle, whereas FuFA-F2 prevented their induction (Figure 7D), supporting that, besides the expression changes observed in several TGF-beta pathway components by RNA-Seq, the anabolic activin-like pathway was suppressed by FuFA-F2. Collectively, these findings link the transcriptional alterations identified in C26 muscle to the activation of multiple pathways involved in muscle wasting, including altered metabolic signaling, autophagy, ubiquitin-dependent proteolysis, and negative regulation of muscle mass. Importantly, FuFA-F2 prevented these alterations, providing molecular evidence that its protective effect against tumor-induced muscle wasting is associated with the preservation of muscle proteostasis.

### 3.7 FuFA-F2 reduced inflammation and fibrosis in skeletal muscle of C26 tumor-bearing mice

Cancer cachexia is frequently associated with systemic and local inflammation [61]. In addition, our RNA-seq analysis revealed a marked alteration of inflammatory and tissue-remodeling pathways in the skeletal muscle of C26-bearing mice, suggesting that tumor growth profoundly affects the local muscle microenvironment. To further characterize these alterations, we investigated systemic and local inflammatory responses as well as muscle fibrosis.

We first assessed circulating inflammation by measuring plasma cytokines using multiplex cytokine analysis. Interestingly, FuFA-F2 supplementation did not significantly modify circulating IL-6 levels in C26- bearing mice, although greater variability was observed in the FuFA-F2-treated group (Figure 8A). Thus, the protective effects of FuFA-F2 on skeletal muscle do not appear to result from a broad suppression of systemic inflammation. We next focused on the local muscle response. Consistent with the inflammatory and tissue-remodeling signatures identified by RNA-seq (e.g. increased expression of key pro-inflammatory pathway components such as Relb, Myd88, Sting, Trex1, Il6, Otd…), *Il6* expression was increased in the muscle of C26 mice, whereas this increase was prevented by FuFA-F2 supplementation (Figure 8B). Similarly, *Cd11b* expression, reflecting the presence of infiltrating myeloid and other immune cells, was increased in C26 muscle but remained at control levels in FuFA-F2-treated mice (Figure 8C). Because inflammatory signaling is closely associated with extracellular matrix remodeling and fibrosis in cachectic muscle, we next assessed collagen deposition by Sirius Red staining. C26 tumor growth was associated with pronounced collagen accumulation in the tibialis muscle, indicative of increased fibrotic remodeling. Notably, FuFA-F2 supplementation significantly attenuated this collagen deposition (Figure 8D). Together, these findings provide functional and histological validation of the inflammatory and tissue-remodeling alterations identified by RNA-seq. Importantly, FuFA-F2 prevented the local inflammatory response, immune cell infiltration, and fibrotic remodeling induced by tumor growth, while having no significant effect on circulating IL-6 levels. These results suggest that FuFA-F2 preserves the local muscle microenvironment rather than exerting a broad systemic anti-inflammatory effect.

**Figure 8.**
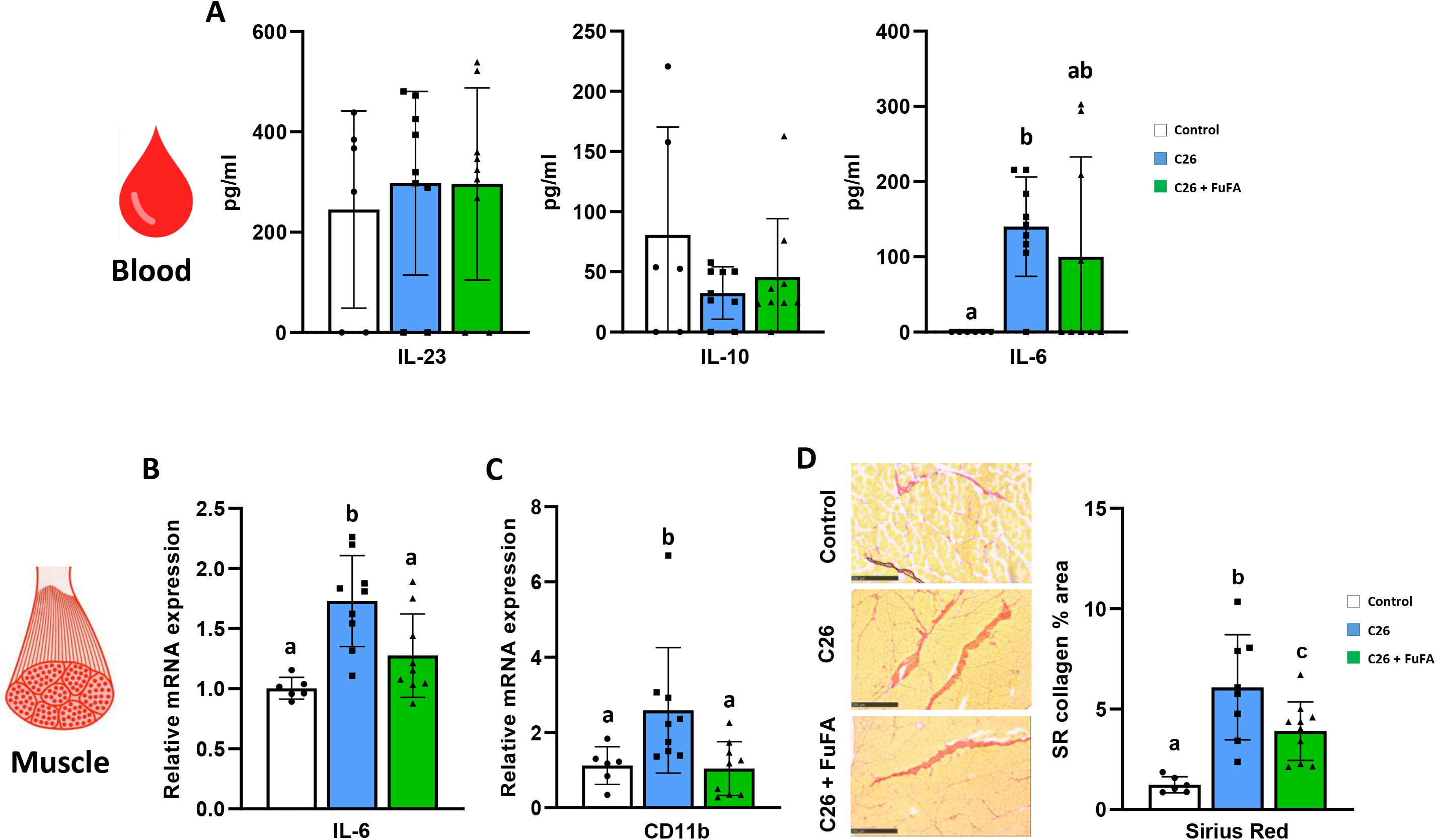
FuFA-F2 reduced fibrosis and inflammation of C26 tumor-bearing mice. (A) Concentration of circulating IL-23, IL-10, and IL-6 in plasma (pg/ml) of the different groups. (B) Relative mRNA expression of *IL-6* in gastrocnemius muscle. (C) Relative mRNA expression of *Cd11b* in gastrocnemius muscle. (D) Representative pictures of transverse cross-sectional area of tibialis muscle after Sirius Red staining. Scale bar, 500µm. Analysis of collagen staining with Sirius Red as a percentage of the area. Results were expressed as means ± SD. Groups were tested by a one-way ANOVA test. The limit of statistical significance was set at p<0.05. The means with different letters were significantly different.

## 4. DISCUSSION

Cancer cachexia is a complex syndrome in which skeletal muscle wasting results from the convergence of metabolic, catabolic, inflammatory, and tissue-remodeling processes. In the present study, we show that FuFA-F2, a naturally occurring furan fatty acid present in small quantities in our diet, markedly protects skeletal muscle from tumor-induced wasting in the C26 model. Importantly, this protective effect occurred despite continued tumor growth and loss of white adipose tissue, indicating that FuFA-F2 does not simply counteract the systemic consequences of tumor progression but exerts a specific protective effect on skeletal muscle. This was supported by both *in vitro* and *in vivo* findings, as FuFA-F2 prevented cytokine-induced myotube atrophy and preserved hindlimb muscle mass in C26-bearing mice. Fatigue, weakness, and reduced physical activity are among the most commonly reported consequences of cancer, which contribute to reduced perceived life quality [56]. In our C26 model, the marked weight loss observed during the final days of the disease was accompanied by a profound decline in spontaneous physical activity, reflecting the severe functional consequences of cancer cachexia. Strikingly, despite comparable tumor progression, C26 mice receiving FuFA-F2 maintained substantially higher levels of voluntary wheel running, with activity levels approaching those of age-matched control mice. We also observed that FuFA-F2-treated animals appeared less prostrate in their cages than untreated C26 mice. These observations suggest that the preservation of skeletal muscle by FuFA-F2 was accompanied by a meaningful functional benefit, allowing tumor-bearing mice to maintain greater physical activity and a more active phenotype despite ongoing disease progression. Thus, beyond preserving muscle mass, FuFA-F2 may contribute to maintaining functional capacity and, at least in part, parameters related to well-being during cancer cachexia.

Our RNA-seq analysis provides important insight into the molecular basis of this protection. Tumor growth induced a profound remodeling of the skeletal muscle transcriptome, with more than 5,400 genes differentially expressed between control and C26 mice. Strikingly, this transcriptional response was markedly attenuated by FuFA-F2, with only a small fraction of genes remaining differentially expressed between control and C26+FuFA-F2 mice. PCA further showed that the transcriptomic profile of FuFA-F2- treated C26 muscle remained close to that of control muscle. Moreover, the strong overlap between genes altered by tumor growth and those responsive to FuFA-F2 suggests that FuFA-F2 predominantly counteracts the transcriptional program induced by the tumor rather than inducing an independent transcriptional state. Together, these findings indicate that FuFA-F2 largely prevents tumor-induced transcriptional reprogramming and preserves a transcriptomic profile closer to that of control skeletal muscle. The nature of the KEGG pathways affected by tumor growth further highlights the broad impact of the tumor on skeletal muscle biology. In addition, C26 muscle showed alterations in pathways involved in protein homeostasis and degradation, including the proteasome, autophagy, and mitophagy. Metabolic and growth-related signaling pathways, such as AMPK, mTOR, and FoxO signaling, were also affected, together with pathways involved in cellular communication and tissue remodeling, including TGF-β signaling. The NF-κB signaling pathway, together with alterations in ribosomal and ribosome biogenesis pathways, further indicates that tumor growth affects both cellular signaling and the machinery supporting protein synthesis. Thus, the transcriptional response to tumor growth was not restricted to a single catabolic pathway but involved multiple interconnected processes regulating protein turnover, cellular signaling, metabolism, and tissue homeostasis. Remarkably, these transcriptional alterations were largely absent or markedly attenuated in C26 + FuFA-F2 muscle when compared with control muscle. This broad preservation of KEGG pathway activity is consistent with the PCA and the limited number of DEGs observed between control and C26 + FuFA-F2 mice, and further supports the concept that FuFA-F2 prevents the extensive transcriptional remodeling induced by tumor growth rather than selectively targeting a single molecular pathway.

The transcriptomic findings were further supported by targeted molecular analyses. C26 tumor growth was associated with increased AMPK activation and induction of the muscle-specific E3 ubiquitin ligases MuRF1 and MAFbx, together with increased expression of the negative regulators of muscle mass Myostatin and Activin A. FuFA-F2 prevented these alterations, providing independent molecular validation of the RNA-seq results. These findings suggest that preservation of muscle mass by FuFA-F2 is associated with maintenance of muscle protein homeostasis and prevention of the activation of several major catabolic pathways. Interestingly, the changes in LC3 and p62 did not follow the classical pattern generally associated with increased autophagy. LC3 levels were reduced and p62 levels increased in C26 muscle, whereas both alterations were prevented by FuFA-F2. Given the advanced stage of cachexia at the time of tissue collection, these findings may reflect profound alterations in the autophagic machinery. However, LC3 and p62 abundance alone cannot establish changes in autophagic flux, and further experiments will be required to determine the precise contribution of autophagy to the protective effect of FuFA-F2.

Our data also indicate that tumor-induced muscle wasting is accompanied by profound remodeling of the local muscle environment. C26 muscle displayed increased expression of inflammatory markers, enhanced immune cell infiltration, and pronounced collagen deposition, indicating the development of a local inflammatory and fibrotic response. Remarkably, FuFA-F2 prevented the increase in muscle *Il6* and *Cd11b* expression, and attenuated collagen accumulation. In contrast, FuFA-F2 did not significantly modify circulating IL-6 levels. These observations suggest that FuFA-F2 does not exert a broad systemic anti-inflammatory effect but rather preserves the local muscle environment. The preservation of the local muscle environment may be particularly relevant to the maintenance of muscle function during cachexia. Chronic inflammation and fibrosis can impair muscle architecture, contractile properties, and tissue regeneration, thereby contributing to functional decline beyond the loss of muscle mass itself. Thus, the ability of FuFA- F2 to limit inflammatory infiltration and fibrotic remodeling, together with its effects on muscle proteostasis and transcriptional regulation, may explain why FuFA-F2-treated C26 mice maintained substantially higher spontaneous physical activity despite ongoing tumor progression.

Taken together, our findings support a model in which FuFA-F2 acts primarily as a muscle-preserving agent during cancer cachexia. FuFA-F2 appears to protect skeletal muscle from the coordinated transcriptional, metabolic, catabolic, inflammatory, and fibrotic responses induced by the tumor. This interpretation is consistent with the marked overlap between tumor-induced and FuFA-F2-responsive transcriptional changes and with the preservation of a transcriptomic profile closer to that of control muscle.

In conclusion, our study demonstrates that FuFA-F2 markedly protects against skeletal muscle wasting in the C26 model of cancer cachexia. FuFA-F2 preserved muscle mass and spontaneous physical activity despite ongoing tumor growth and adipose tissue loss, while largely preventing the extensive transcriptional reprogramming induced by the tumor. This protection was associated with preservation of muscle proteostasis and reduced catabolic, inflammatory, and fibrotic remodeling. Thus, FuFA-F2 appears to act primarily by preserving skeletal muscle integrity and its local environment rather than by suppressing tumor growth or systemic inflammation. These findings identify FuFA-F2 as a promising candidate for therapeutic strategies aimed at preserving skeletal muscle during cancer cachexia.

## ACKNOWLEDGMENTS

The authors also wish to thank the Metamus DMeM facility (https://doi.org/10.15454/WYR2-8706), which belongs to the Montpellier animal facilities network (RAM, BioCampus) for technical support and expertise for metabolism phenotyping. The authors also thank the “Réseau d’Histologie Expérimentale de Montpellier” (RHEM, BioCampus) and “Montpellier Ressources Imagerie”(MRI, BioCampus).

## DATA AVAILABILITY STATEMENT

All data are available on request.

## HIGHLIGHTS

- FuFA-F2 prevented muscle wasting in C26 tumor-bearing mice.
- FuFA-F2 markedly attenuates tumor-induced muscle transcriptional reprogramming.
- FuFA-F2 preserves muscle proteostasis and limits catabolic pathways.
- FuFA-F2 attenuates local muscle inflammation and fibrotic remodeling.
- FuFA-F2 might be a promising approach to counteract cancer cachexia–induced muscle wasting.

## AUTHOR CONTRIBUTIONS

A.D., J.F., C.G., C.B-G., B.B., V.B., L.H-M., E.D., P.D., C.J-T., C.D., A.C., J.C., L.V., S.L., L.P., and F.C acquisition of data. A.D., C.G., C.B-G., V.B., L.H-M., E.D., C.C., W.V, C.F-C., A.D. and F.C. analysis and interpretation of data. A.D. and F.C. conception and design, wrote the paper. All the authors have read and approved the final version of this manuscript.

---

**Supplemental Figure 1.** Primers used in Q-PCR experiments.

**Supplemental Figure 2.** Comparison of KEGG Pathway of Proteasome with visualization of gene expression differences between Control vs C26 and C26 vs C26+FuFA. The genes boxed in purple are the DEGs.

**Supplemental Figure 3.** Comparison of KEGG Pathway of Autophagy with visualization of gene expression differences between Control vs C26 and C26 vs C26+FuFA. The genes boxed in purple are the DEGs.

**Supplemental Figure 4.** Comparison of KEGG Pathway of Mitophagy in cancer with visualization of gene expression differences between Control vs C26 and C26 vs C26+FuFA. The genes boxed in purple are the DEGs.

**Supplemental Figure 5.** Comparison of KEGG Pathway of AMPK signaling pathway with visualization of gene expression differences between Control vs C26 and C26 vs C26+FuFA. The genes boxed in purple are the DEGs.

**Supplemental Figure 6.** Comparison of KEGG Pathway of TGF-BETA signaling pathway with visualization of gene expression differences between Control vs C26 and C26 vs C26+FuFA. The genes boxed in purple are the DEGs.

**Supplemental Figure 7.** Comparison of KEGG Pathway of FOXO signaling pathway with visualization of gene expression differences between Control vs C26 and C26 vs C26+FuFA. The genes boxed in purple are the DEGs.

**Supplemental Figure 8.** Comparison of KEGG Pathway of NF-KAPPA B signaling pathway with visualization of gene expression differences between Control vs C26 and C26 vs C26+FuFA. The genes boxed in purple are the DEGs.

**Supplemental Figure 9.** Comparison of KEGG Pathway of Ribosomes Biogenesis with visualization of gene expression differences between Control vs C26 and C26 vs C26+FuFA. The genes boxed in purple are the DEGs.

**Supplemental Figure 10.** Comparison of KEGG Pathway of Ribosomes with visualization of gene expression differences between Control vs C26 and C26 vs C26+FuFA. The genes boxed in purple are the DEGs.

**Supplemental Figure 11.** Comparison of KEGG Pathway of mTOR signaling pathway with visualization of gene expression differences between Control vs C26 and C26 vs C26+FuFA. The genes boxed in purple are the DEGs.

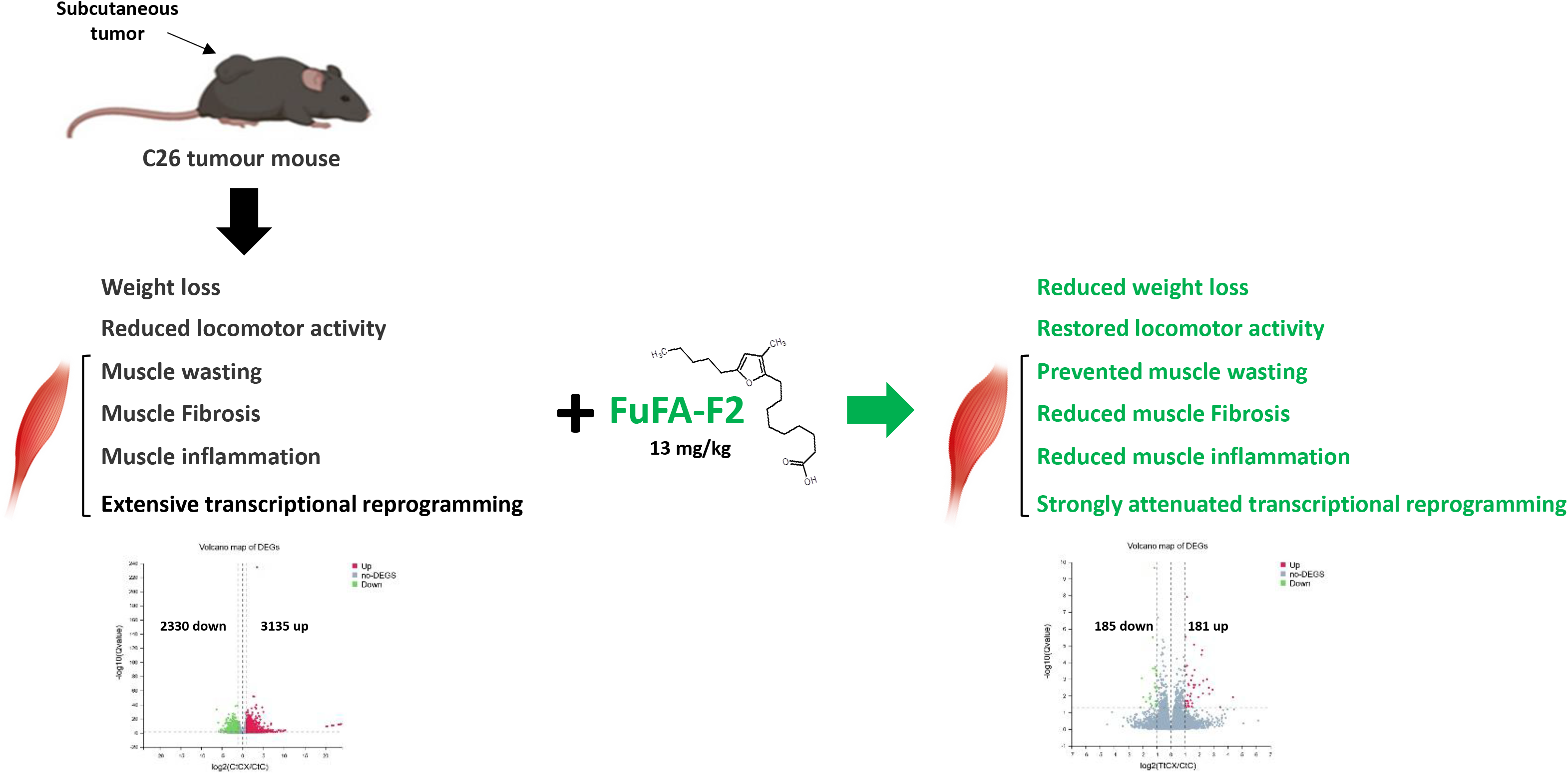

