## Supplemental figure 1-11 for "Furan fatty acid supplementation protects against muscle atrophy during cancer cachexia": Supplemental fig 1-11.pdf

| | | |
|---|---|---|
| Act-A | GGCTGGCATACCTTTCCCAT | GTTTGCGGATGCGATGTCTG |
| Mstn | GCACTGGTATTTGGCAGAGTA | CACACTCTCCTGAGCAGTAAT |
| Fbxo32 (MAFbx) | TCAGAGAGGCAGATTCGCAA | GGGTGACCCCATACTGCTCT |
| Trim63 (Murf1) | TCCTGATGGAAACGCTATGGAG | ATTCGCAGCCTGGAAGATGT |
| IL6 | AAGACAAAGCCAGAGTCCTTCA | GCATTGGAAATTGGGGTAGGAAG |
| CD11b | TGTGGACTCTCATGCCTCCT | TGGTCATCTCTGAAGCCGTG |

**Supplemental Figure 1**

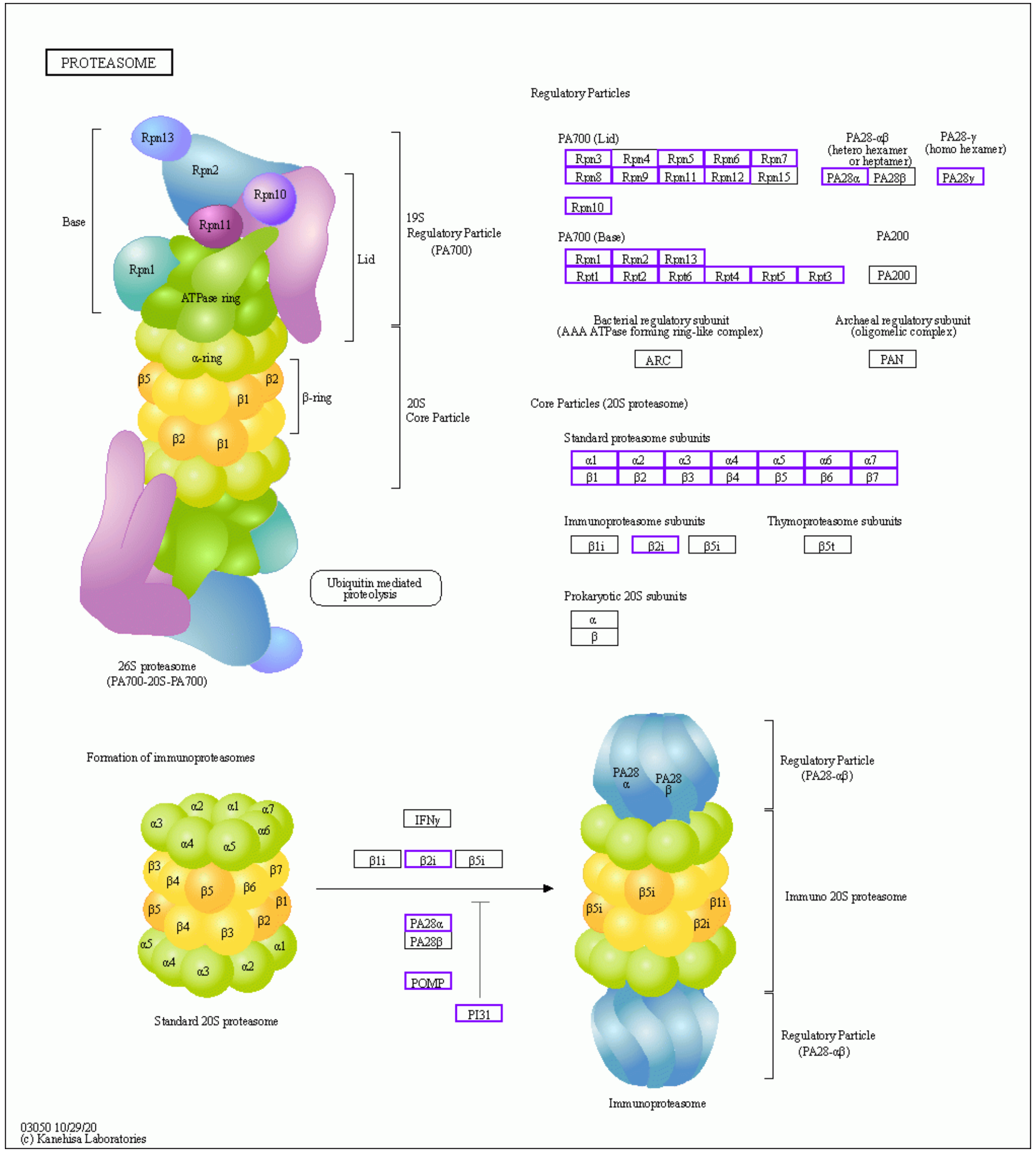


Control vs C26

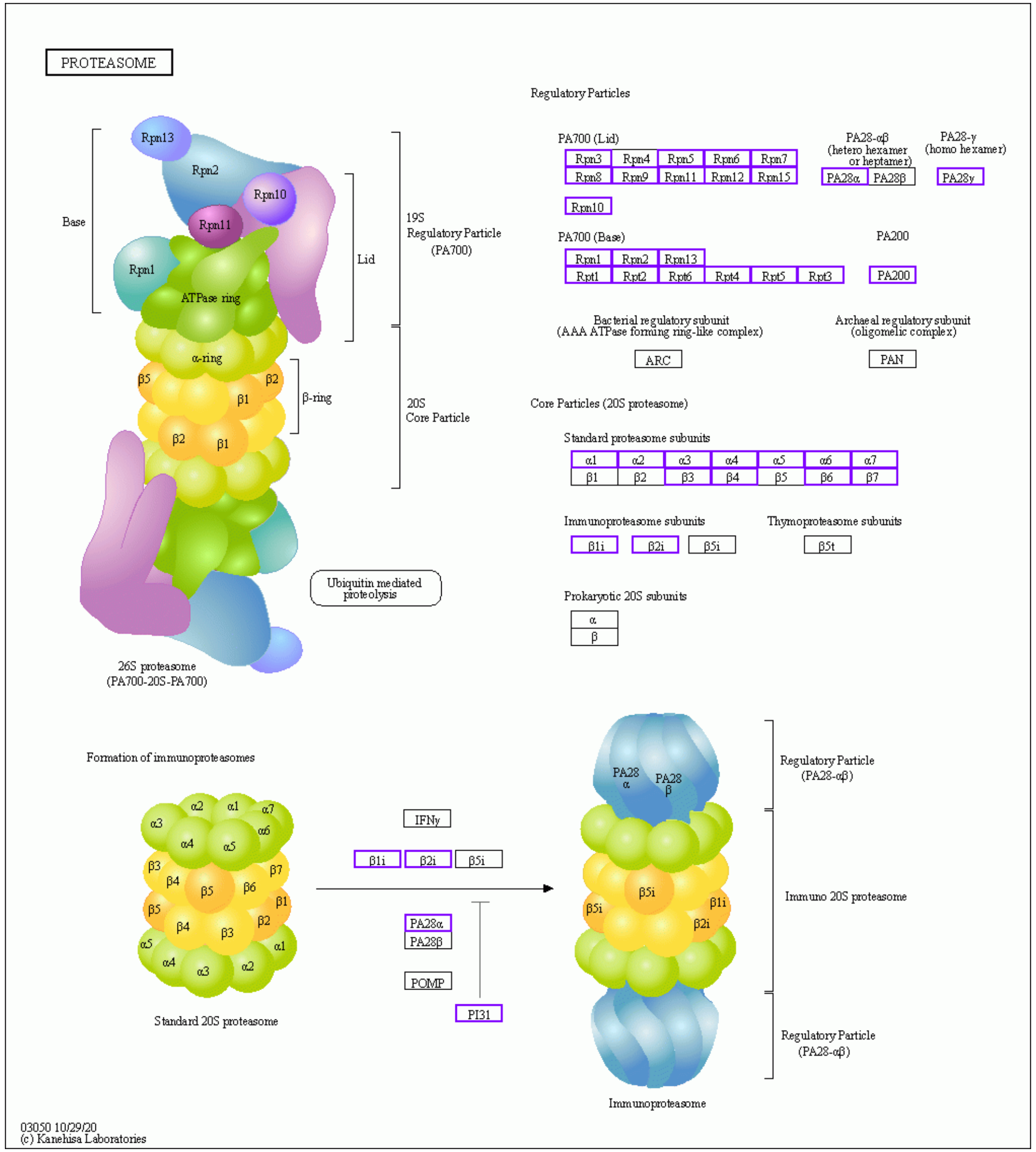


C26 vs C26+FuFA

**Supplemental Figure 2**

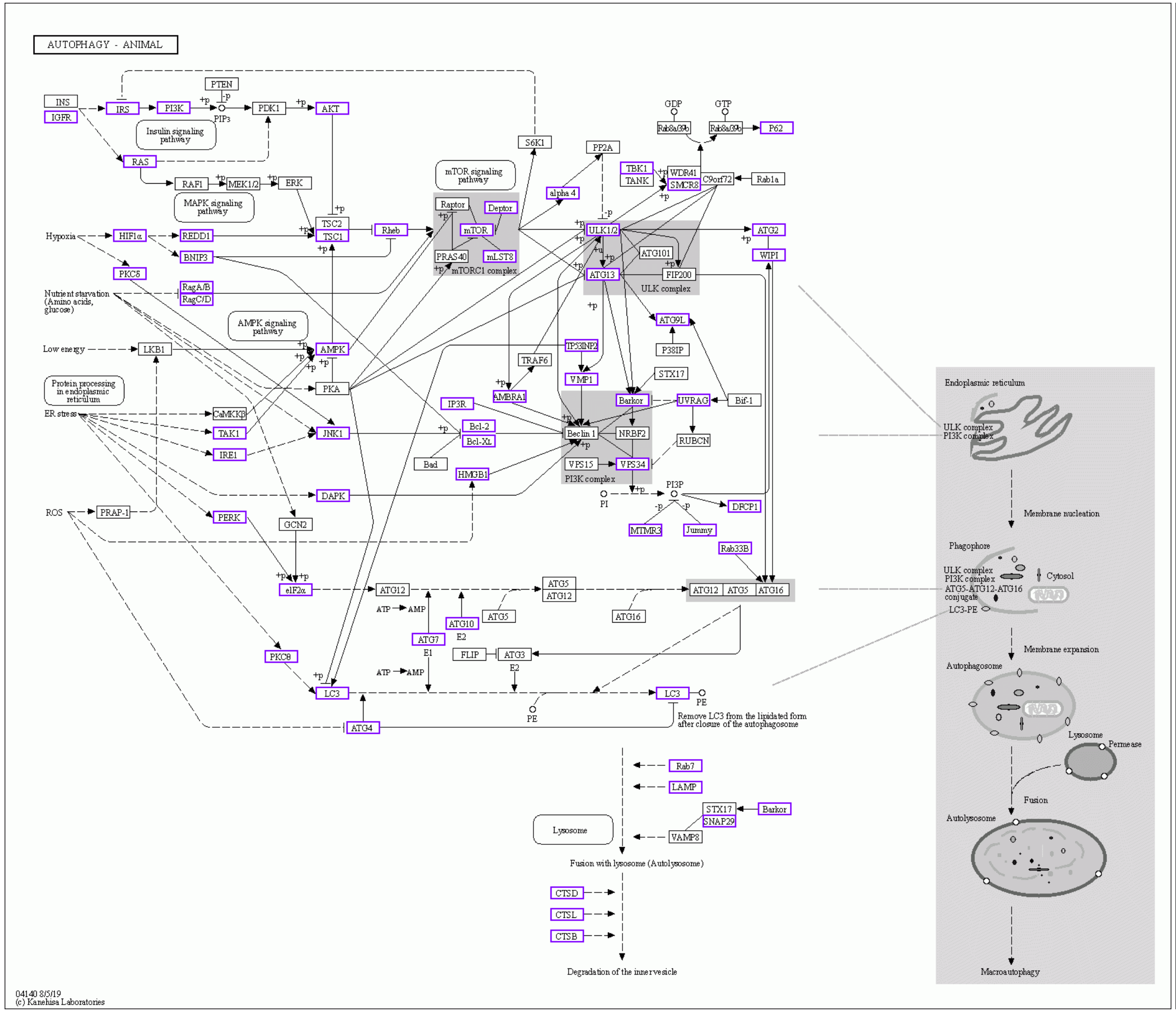


Control vs C26

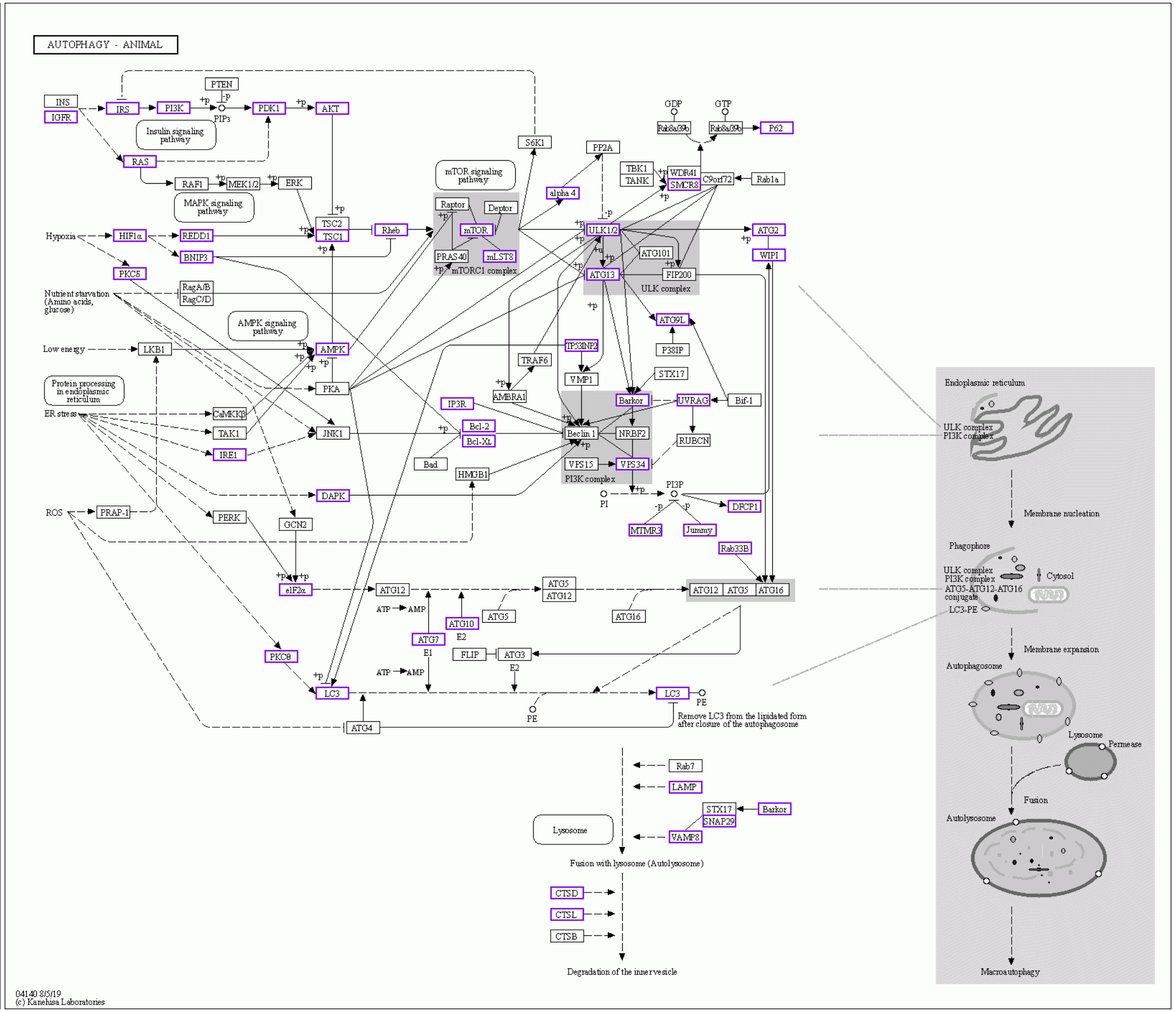


C26 vs C26+FuFA

**Supplemental Figure 3**

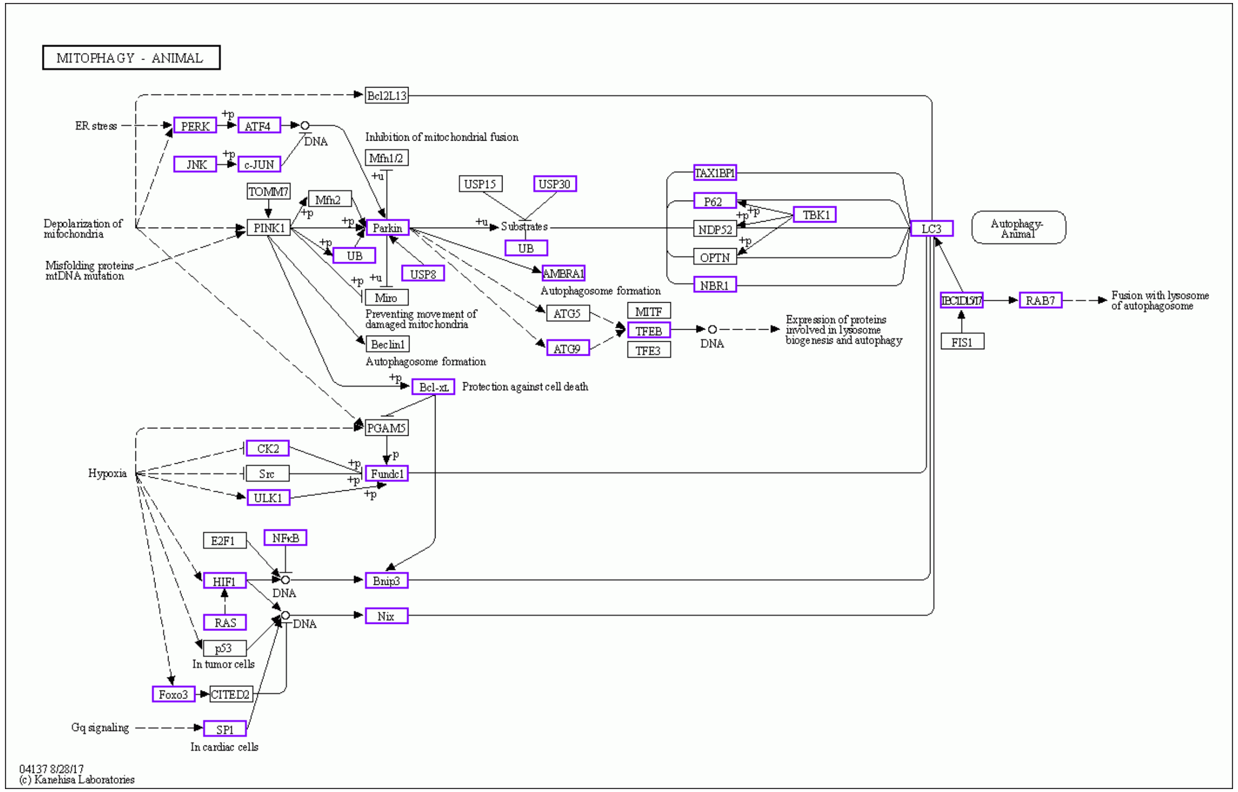


**Control vs C26**

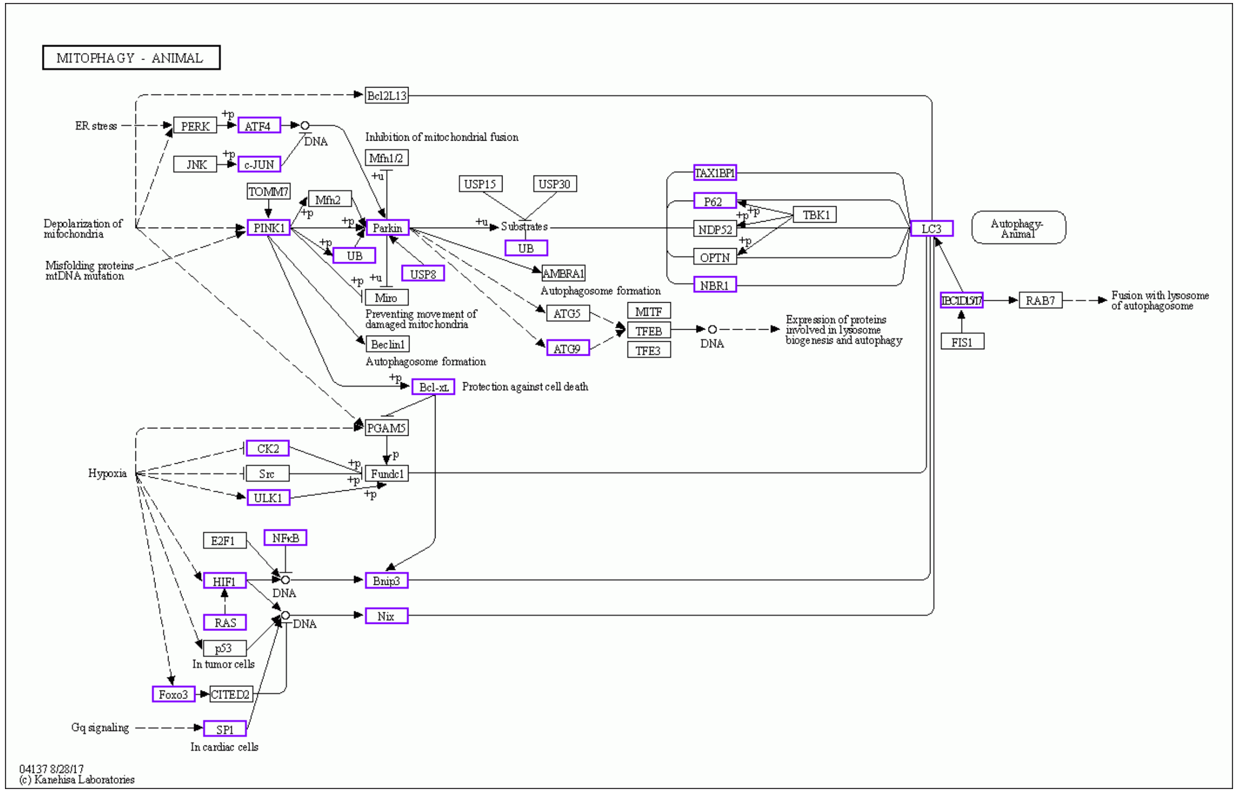


**C26 vs C26+FuFA**

**Supplemental Figure 4**

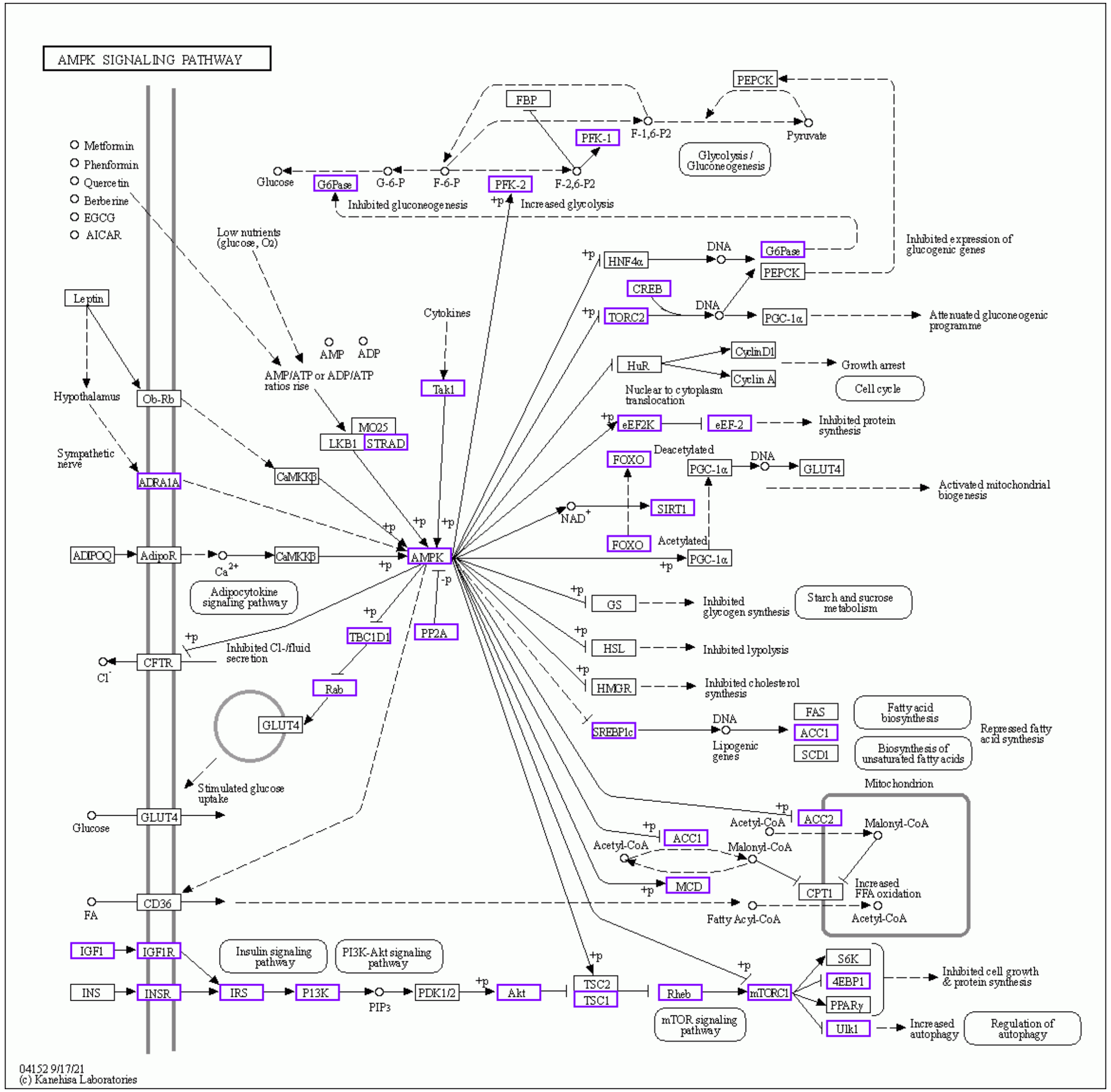


Control vs C26

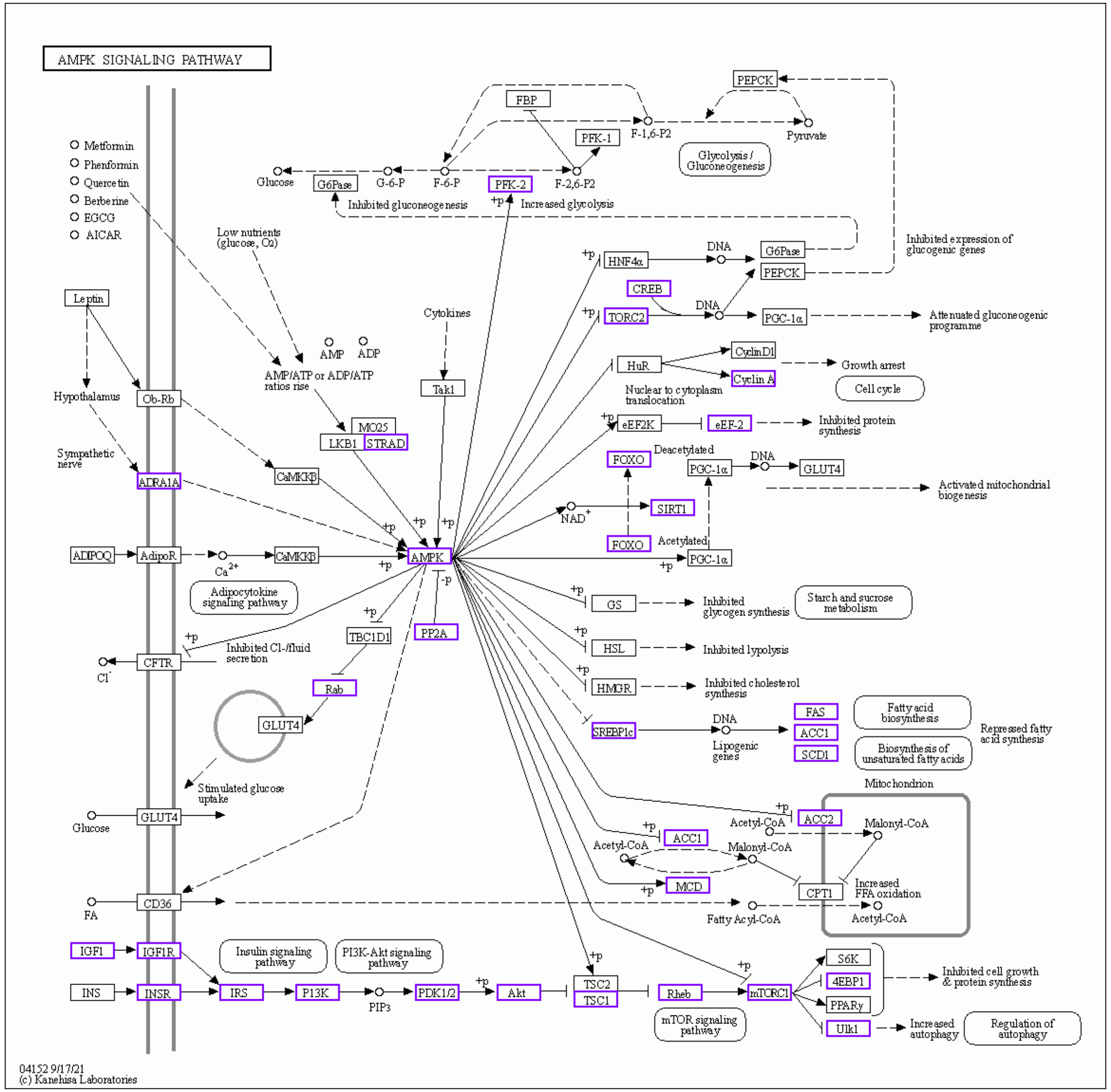


C26 vs C26+FuFA

**Supplemental Figure 5**

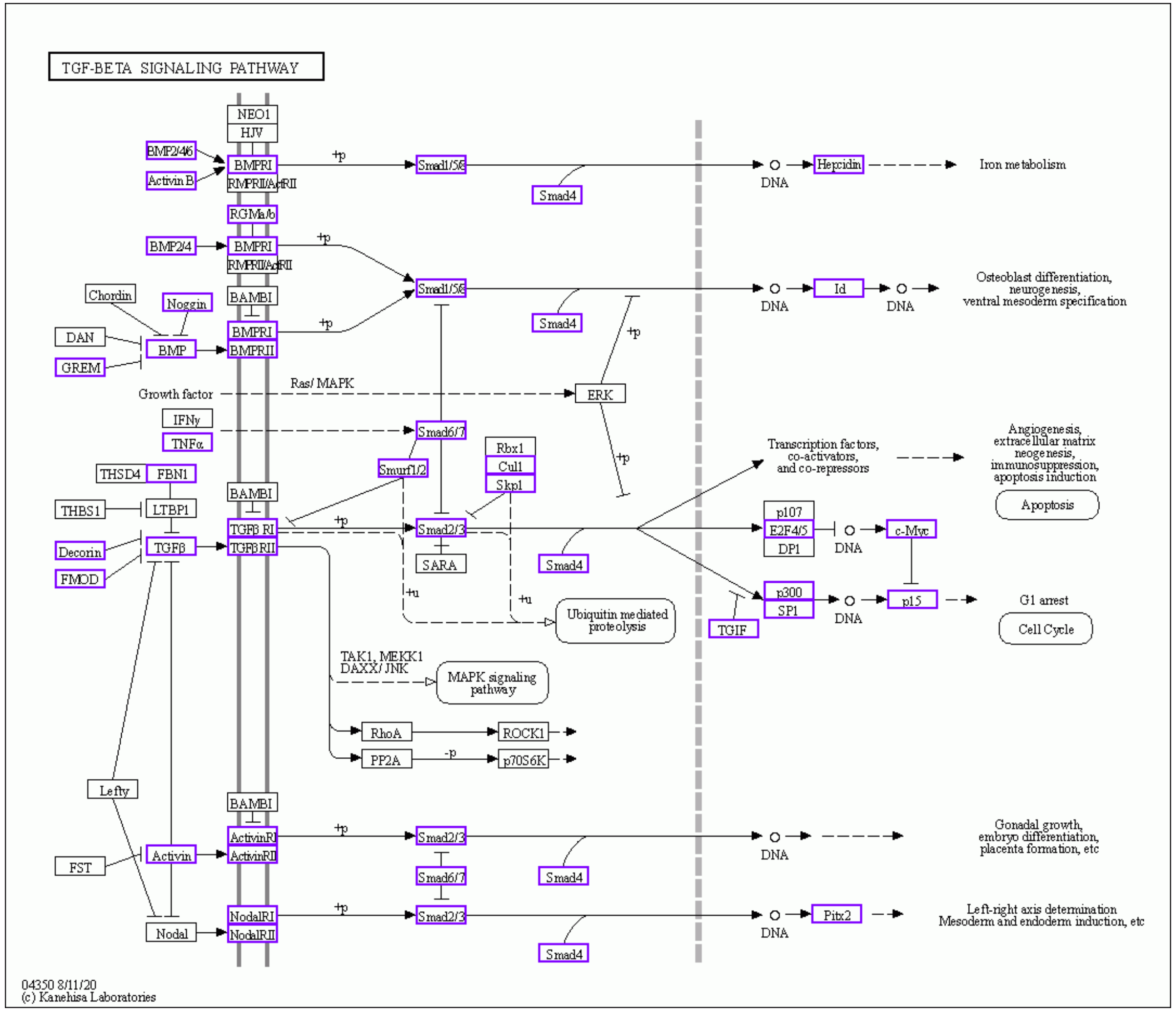


Control vs C26

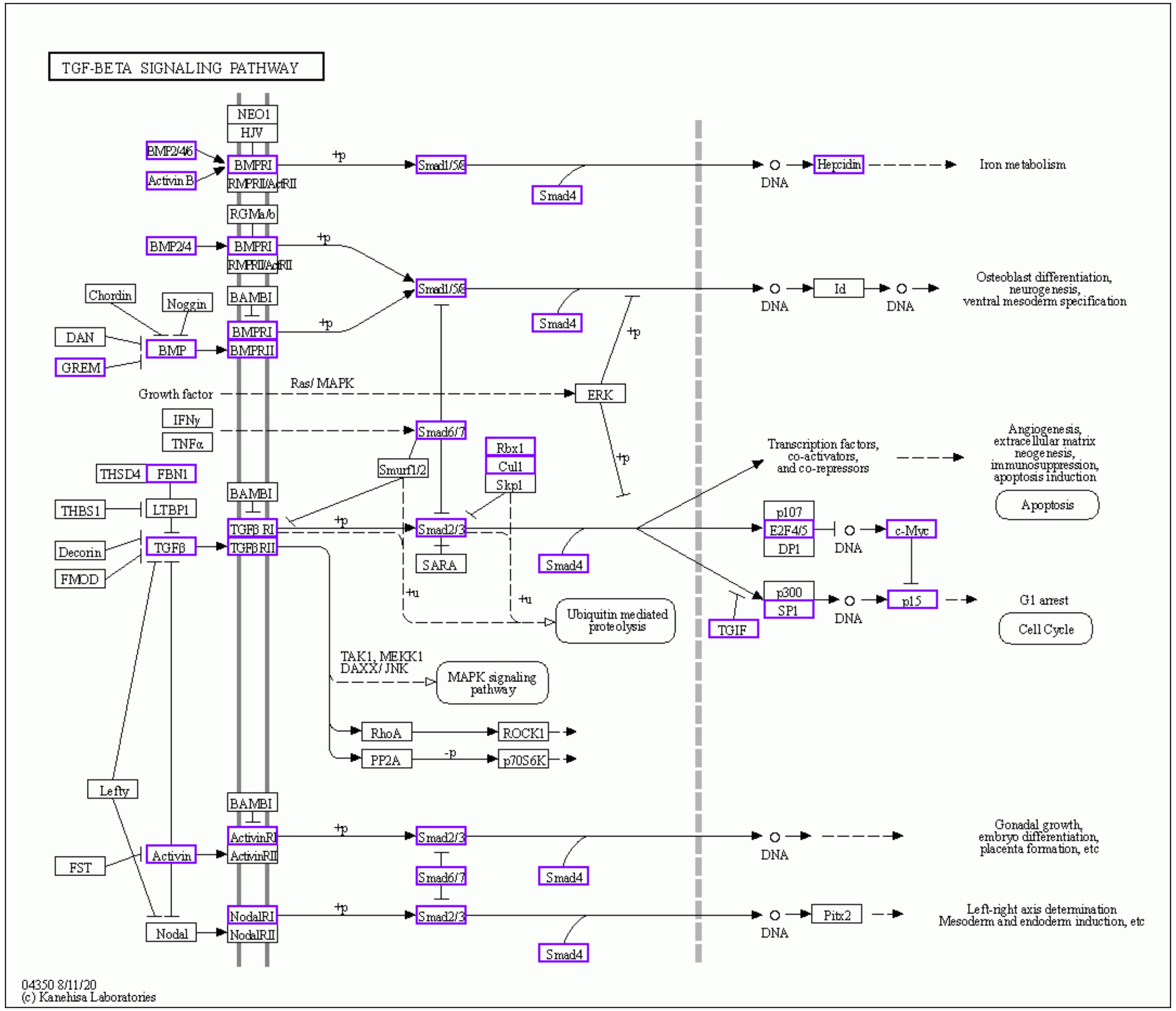


C26 vs C26+FuFA

**Supplemental Figure 6**

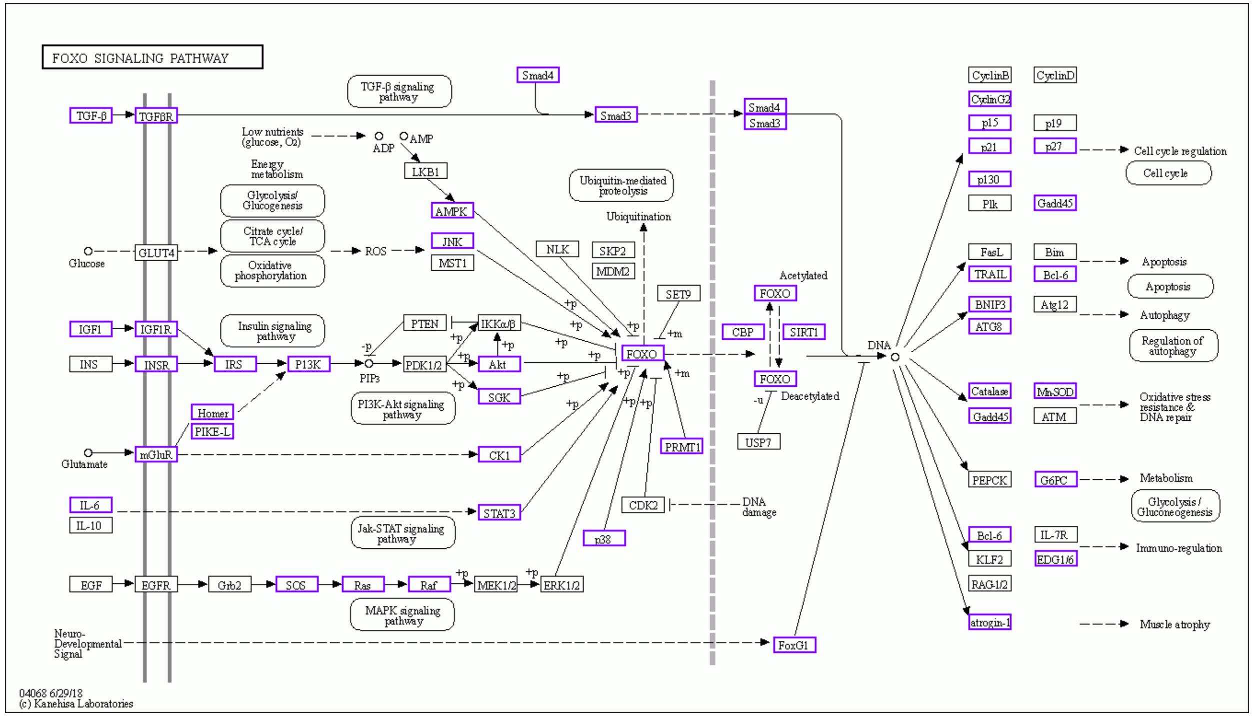


**Control vs C26**

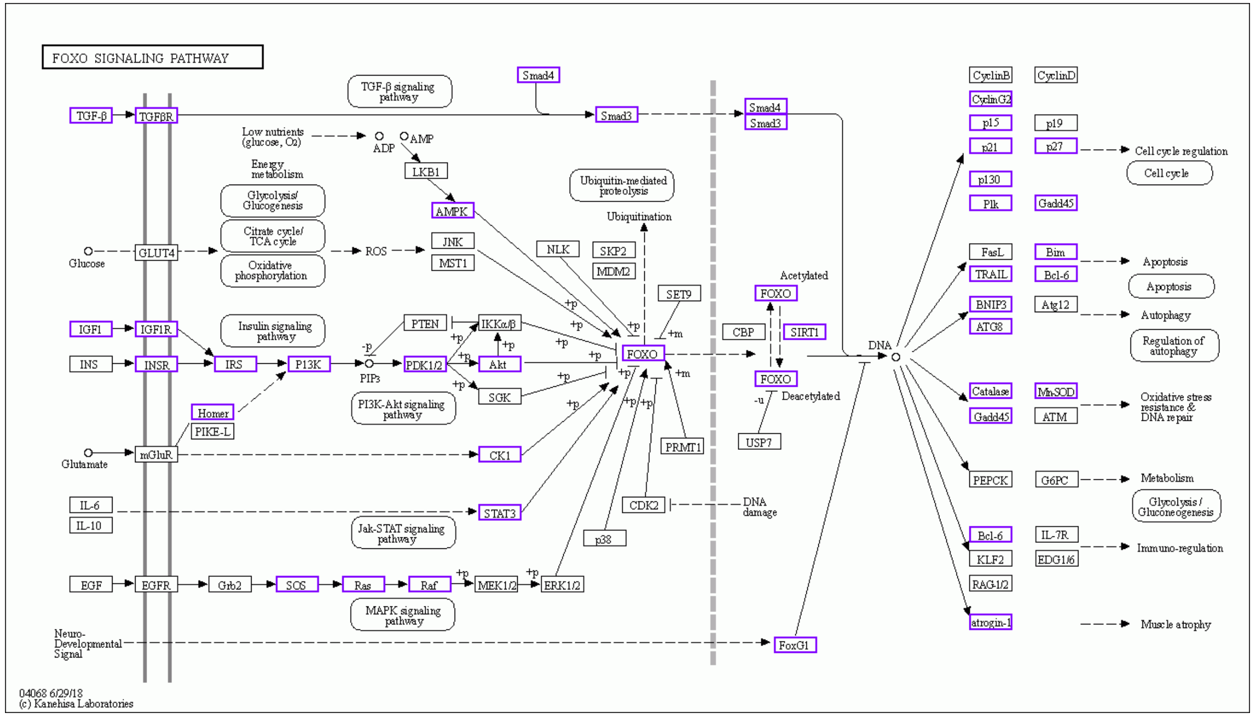


**C26 vs C26+FuFA**

### Supplemental Figure 7

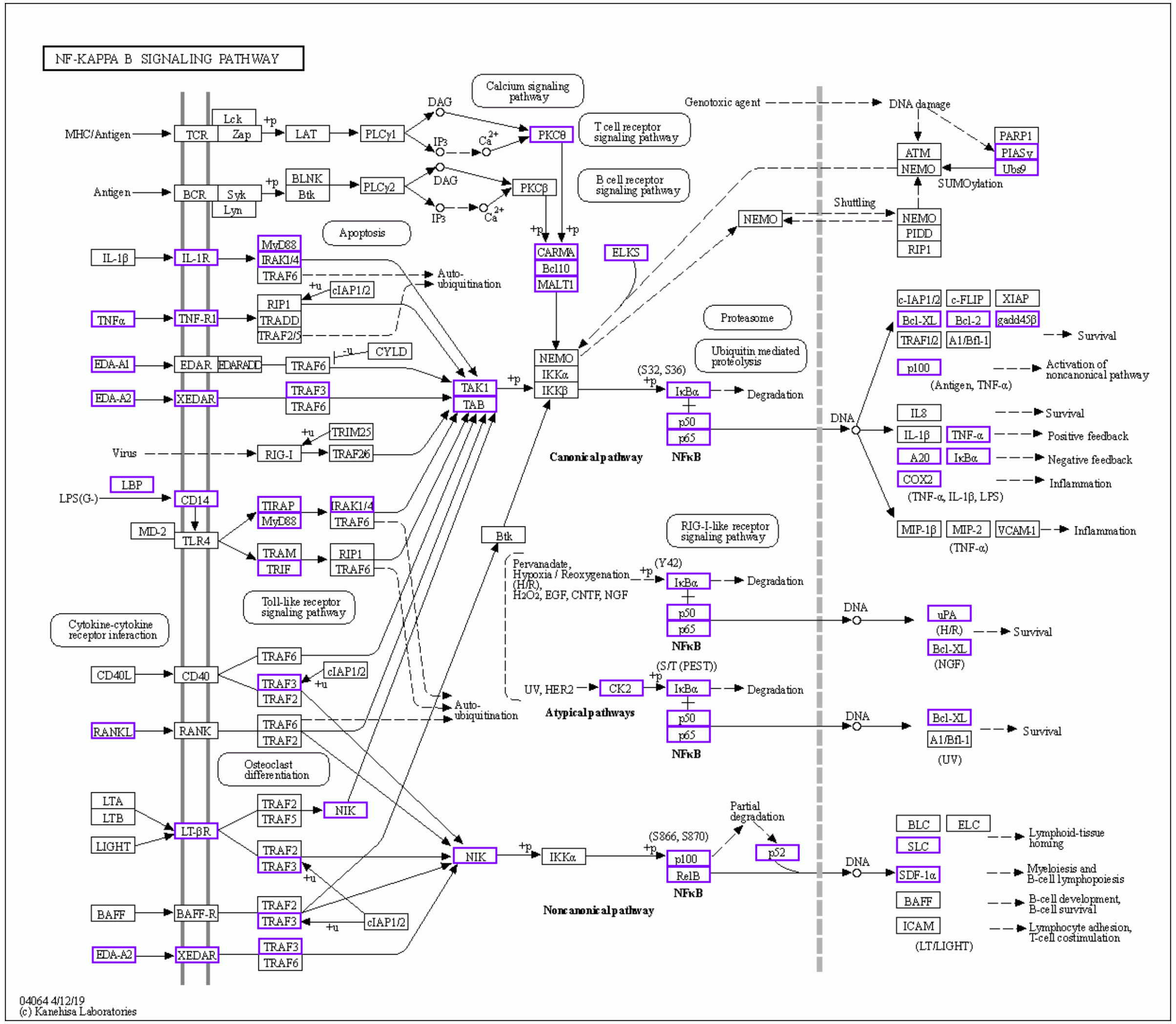


Control vs C26

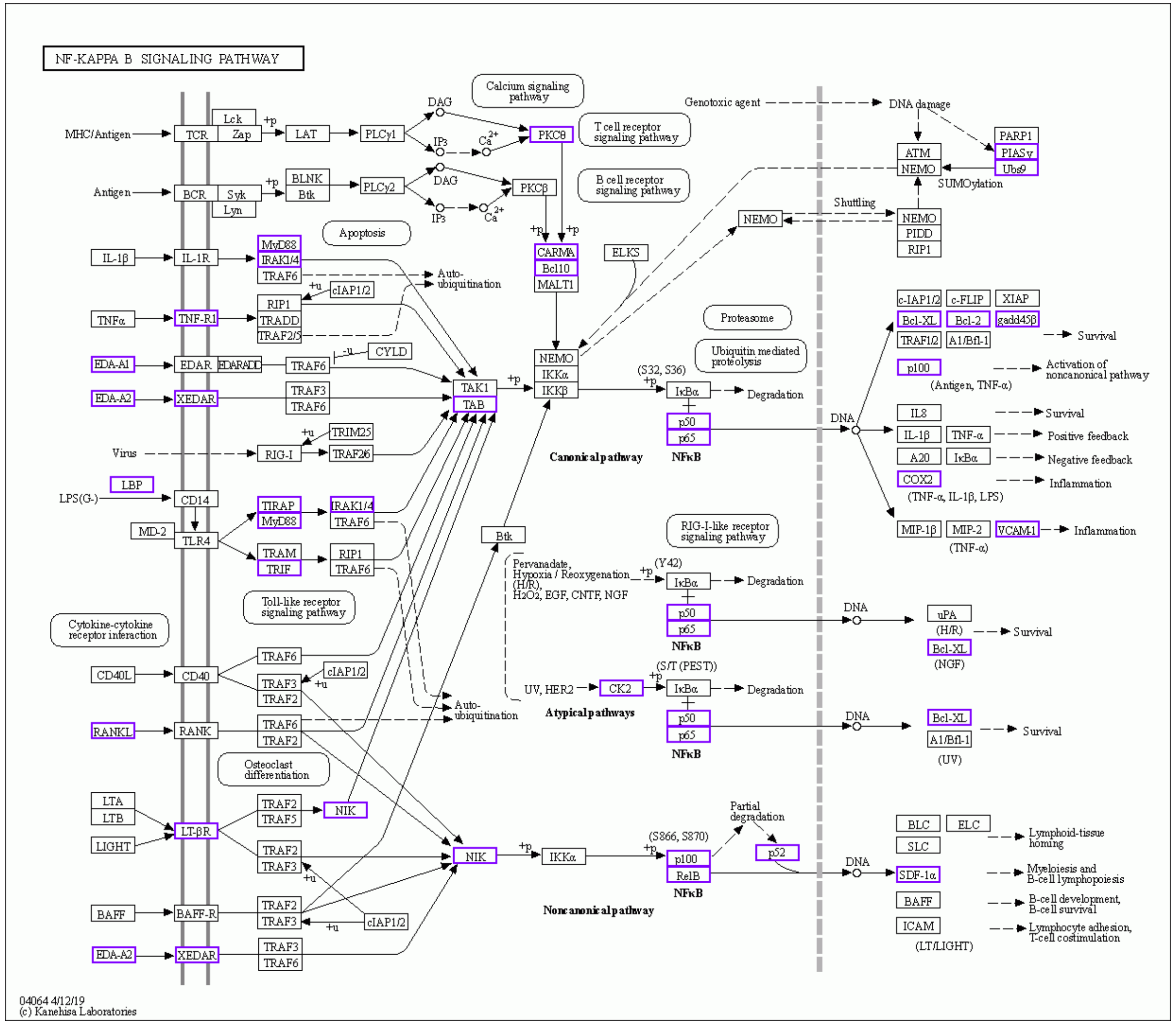


C26 vs C26+FuFA

**Supplemental Figure 8**

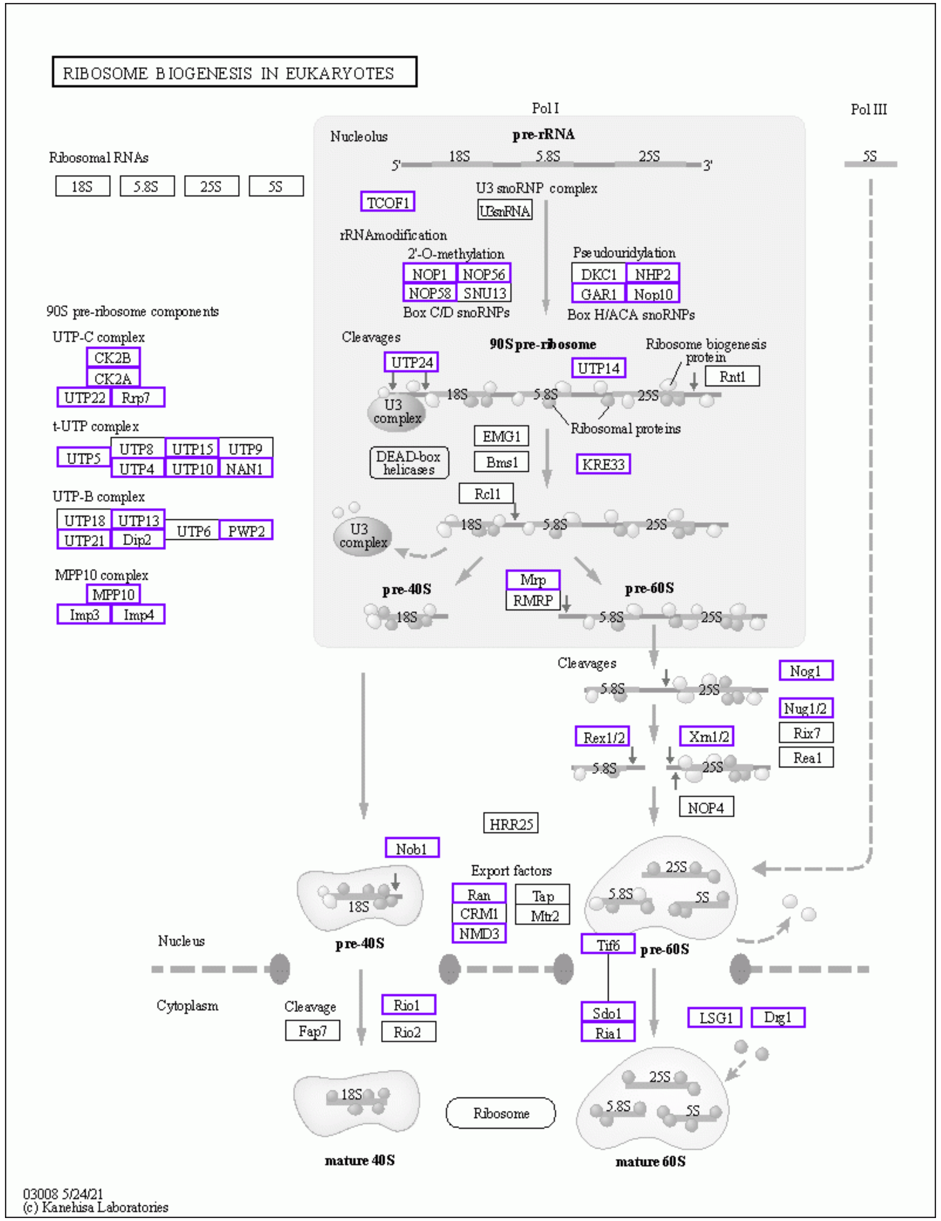


**Control vs C26**

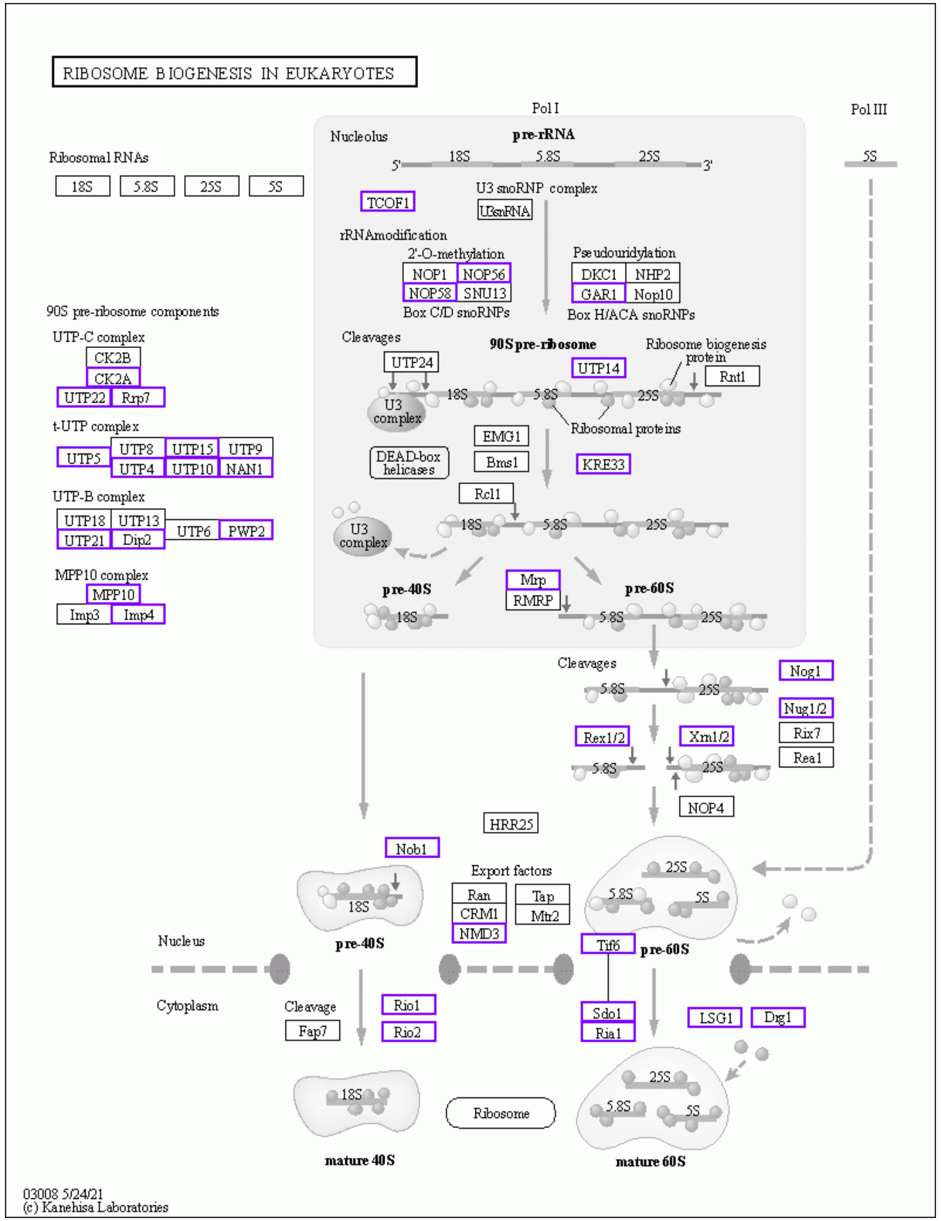


**C26 vs C26+FuFA**

**Supplemental Figure 9**

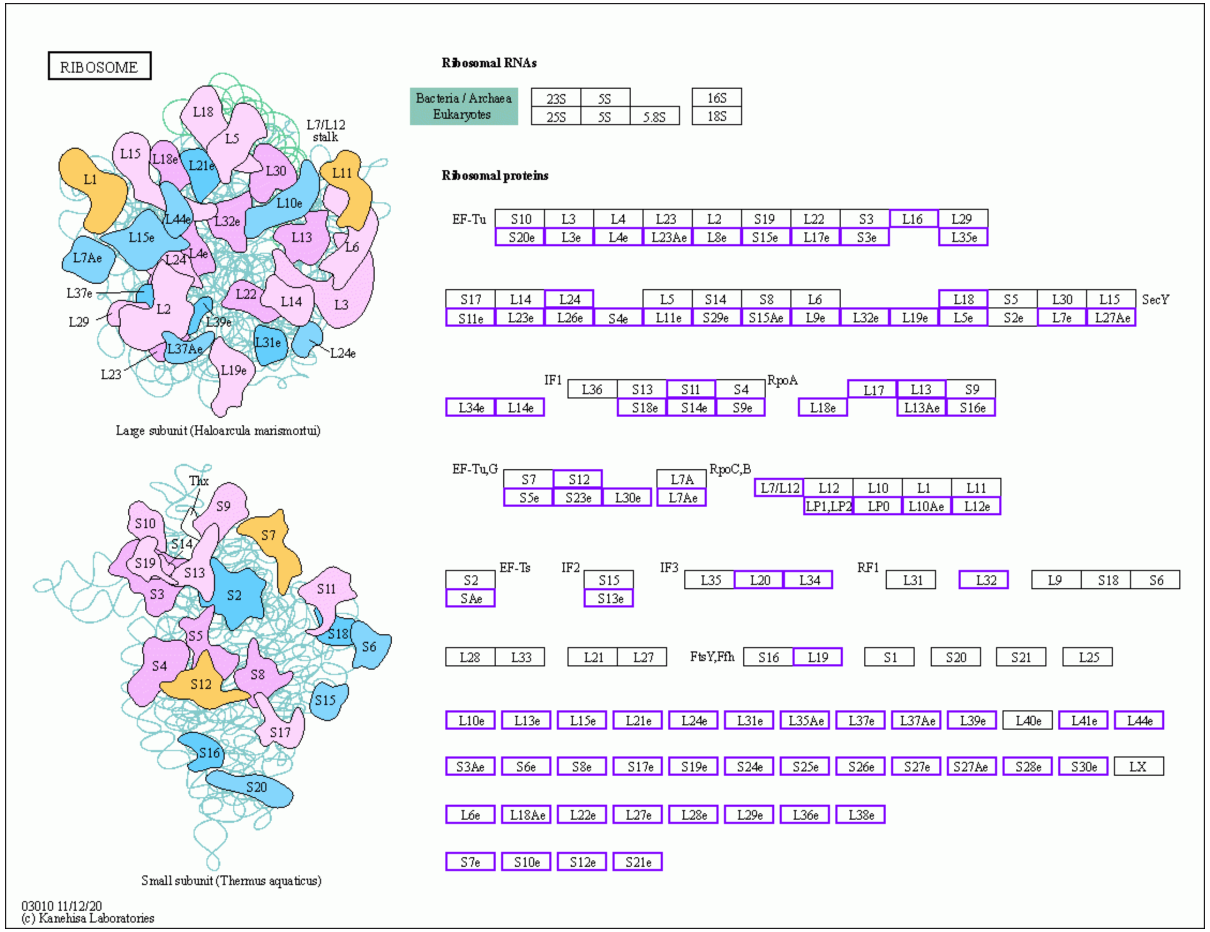


Control vs C26

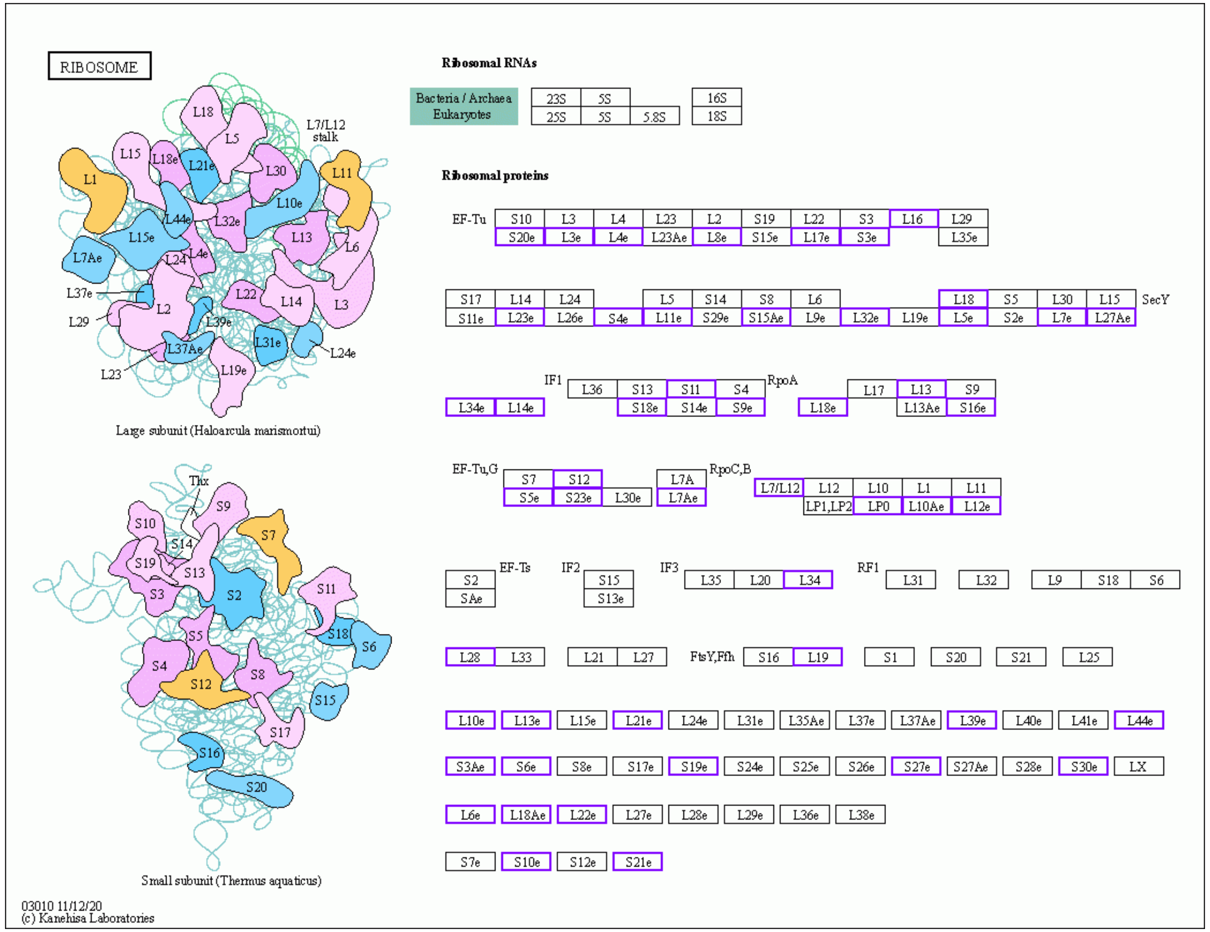


C26 vs C26+FuFA

**Supplemental Figure 10**

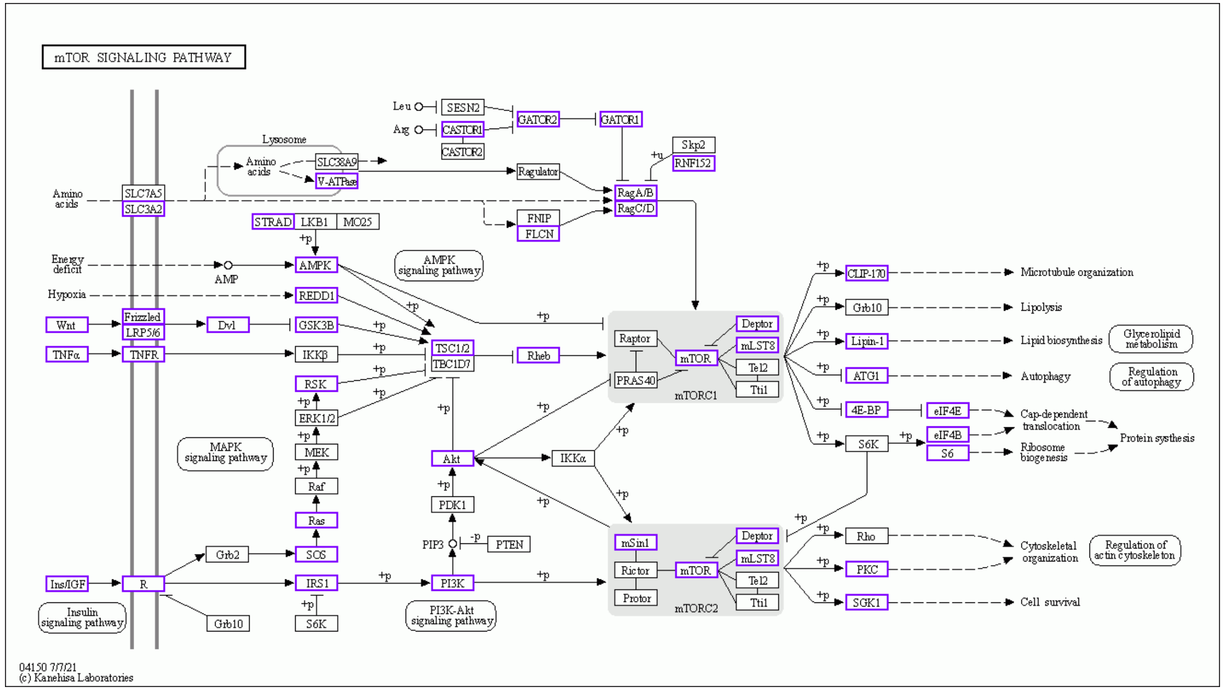


**Control vs C26**

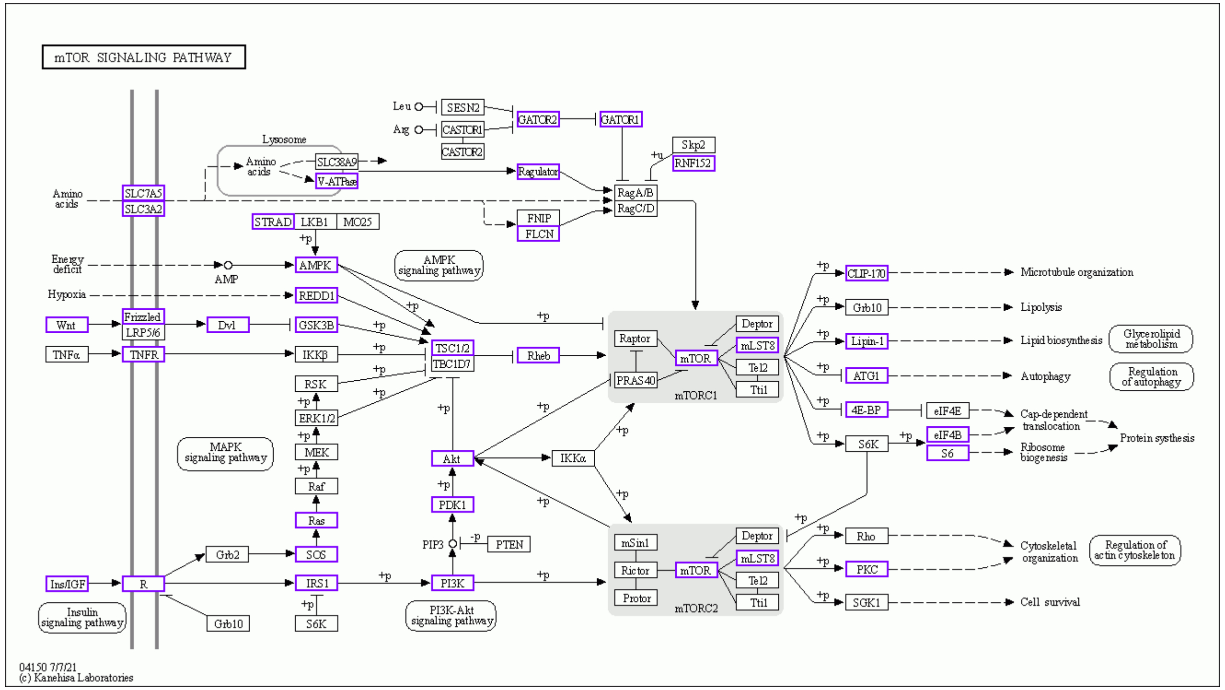


**C26 vs C26+FuFA**

### Supplemental Figure 11
